# βA3/A1-crystallin is an epigenetic regulator of histone deacetylase 3 (HDAC3) in the retinal pigmented epithelial (RPE) cells

**DOI:** 10.1101/2024.08.06.606634

**Authors:** Sujan Chatterjee, Sayan Ghosh, Zachary Sin, Emily Davis, Loretta Viera Preval, Nguyen Tran, Sridhar Bammidi, Pooja Gautam, Stacey Hose, Yuri Sergeev, Miguel Flores-Bellver, Issam Aldiri, Debasish Sinha, Prasun Guha

## Abstract

The retinal pigmented epithelial (RPE) cells maintain retinal homeostasis, and alterations in their function contribute to non-exudative age-related macular degeneration (AMD)^1,2^. Here, we explore the intricate relationship between RPE cells, epigenetic modifications, and the development of AMD. Importantly, the study reveals a substantial decrease in histone deacetylase 3 (HDAC3) activity and elevated histone acetylation in the RPE of human AMD donor eyes. To investigate epigenetic mechanisms in AMD development, we used a mouse model with RPE-specific *Cryba1* knockout^3–5^, revealing that the loss of βA3/A1-crystallin selectively reduces HDAC3 activity, resulting in increased histone acetylation. βA3/A1-crystallin activates HDAC3 by facilitating its interaction with the casein kinase II (CK2) and phosphorylating HDAC3, as well as by regulating intracellular InsP6 (phytic acid) levels, required for activating HDAC3. These findings highlight a novel function of βA3/A1-crystallin as an epigenetic regulator of HDAC3 in the RPE cells and provide insights into potential therapeutic strategies in non-exudative AMD.

## Main Text

The retinal pigmented epithelial (RPE) cells serve as specialized phagocytes of the eye and persevere throughout an organism’s entire lifespan^1^. These post-mitotic cells play a vital role in maintaining the homeostasis of the neural retina, a function essential for normal vision^1^. In recognizing the pivotal role of RPE in ocular health, it has become evident that alterations in RPE function can trigger the atrophic form of age-related macular degeneration (AMD)^1–6^, a debilitating eye disease prevalent in the aging population with limited therapeutic options. Despite Genome-Wide Association Studies (GWAS) identifying several AMD risk variants, their effects remain modest, leaving approximately 70% of AMD risk unexplained. Notably, early-stage AMD is associated with decreased chromatin accessibility in the RPE^6^, suggesting a potential link between epigenetic changes in RPE cells and disease onset. This highlights the notion that, in a multifactorial aging disease like AMD, epigenetic factors may exert a substantial influence beyond genetic variations. To gain deeper insights into the identification of epigenetic mechanisms in the RPE influencing the development of AMD, we utilized a mouse model with a specific knockout of *Cryba1*^3–5^, the gene encoding βA3/A1- crystallin, in the RPE cells. This *Cryba1* knockout (cKO) mouse model is a valuable tool for investigating the complex mechanisms underlying AMD^3–5^, as it replicates numerous clinical features and molecular changes observed in human AMD patients^3–5^. Since human atrophic AMD tissue is limited, such models are particularly important. Our research with the *Cryba1* cKO model bridges this gap, providing a crucial link to understanding the impact of epigenetic factors on AMD development and validating these findings in the context of human disease.

In previous studies, it was established that βA3/A1-crystallin plays a vital role in regulating the mechanistic target of rapamycin (mTOR) and lysosomal function^3^. This regulation is achieved by binding with the V0 subunit of V-ATPase, a proton pump essential for lysosome acidification^4^. Loss of βA3/A1-crystallin protein caused reduction in V-ATPase activity and subsequent increase in lysosomal pH, activation of mTORC1, inhibition of autophagy, and disruption of the lysosomal clearance machinery^3^. This study explores the possibility of epigenetic modifications in an AMD-like mouse model, examining how they may influence the control of various physiological activities in RPE cells by βA3/A1-crystallin. RNA-seq analysis revealed that βA3/A1-crystallin deletion significantly impacts global transcription, resulting in upregulating 251 genes and downregulating 290 genes **(Extended Figure 1A)**. Histone acetylation is a reversible epigenetic modification crucial in regulating gene expression^7^. This dynamic process involves the addition of acetyl groups to histone proteins, which can directly influence the accessibility of DNA and, consequently, the activation or repression of specific genes^7^. Interestingly, we observed a marked increase in global H3/H4 acetylation in *Cryba1* cKO RPE cells compared to *Cryba1* floxed (*Cryba1*^fl/fl^) controls, including H3K9, H3K18, H3K27, H4K12 and H4K16 acetylation **(Extended Data** Figs. 2A-J). These effects closely resembled the impact of SAHA (Suberoylanilide Hydroxamic acid) **(Extended Data** Figs. 2J), a pan histone deacetylase inhibitor used as a positive control, which is known for enhancing global histone acetylation^8^.

In cells, histone acetylation levels are controlled by histone “writers” and “erasers.” P300/CBP is a major “writer” that adds acetyl groups to histone lysine, while histone deacetylases (HDACs) act as “erasers” by removing these groups^9^. Our hypothesis suggests two possibilities: i) the loss of *Cryba1* increases the activity of histone acetyltransferase (HAT), or ii) it reduces the activity of HDACs, resulting in a significant rise in histone acetylation. Interestingly, when *Cryba1* is deleted, global class I HDAC activity is notably reduced in the cKO RPE, compared to floxed controls **(Figure 1A, B)**, but HAT activity remains unchanged **(Extended Figure 3A)**. A comprehensive examination of all class I HDACs (HDAC1, 2, 3, and 8) revealed that loss of *Cryba1* selectively diminished HDAC3 activity (**Figure 1F)** (> 82.97%), leaving other class I HDACs unaffected (**Figure 1C-H)**. Intriguingly, *Cryba1* loss did not alter the levels of HDAC3 protein **(Figure 1I)**. Moreover, reintroducing *Cryba1* using an adenoviral vector in cKO RPE cells restored HDAC3 activity to near-normal levels, unlike the control vector **(Extended Figure 3B)**. Additionally, *Cryba1* overexpression reduced histone acetylation levels, particularly for H3K9 **(Extended Figure 3C, E)** and H3K27 **(Extended Figure 3D, F).** These findings highlight the role of *Cryba1* in selectively regulating HDAC3 activity and modulating histone acetylation in RPE cells. *Cryba1* produces two protein isoforms, βA3- and βA1-crystallin, from the same mRNA by leaky ribosomal scanning^10^. It is now recognized that different isoforms resulting from alternative translation pathways may exhibit unique and novel functionalities^10^. Notably, our findings revealed that RPE cells obtained from CRISPR/cas9 gene-edited mice with βA1-crystallin knockdown (KD) had a more pronounced decrease in HDAC3 activity than those with βA3-crystallin knockout (KO); however, the protein levels of HDAC3 remained unchanged in both isoforms **(Figure 1J, K)**.

**Figure 1:**
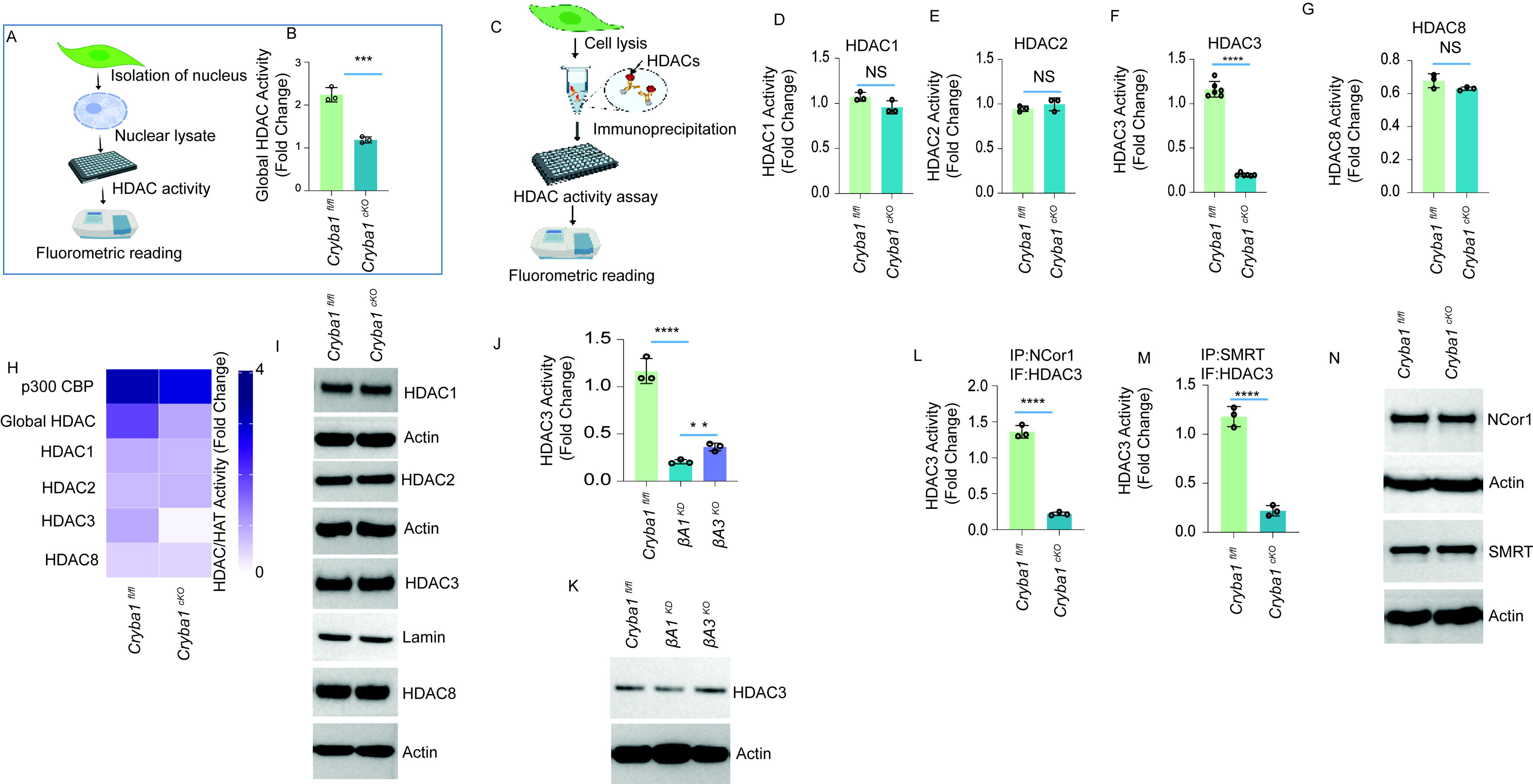
*Cryba1* deletion diminishes HDAC3 activity in mouse RPE cells. **(A)** Schematic diagram of global HDAC activity assay. Nuclear lysate was isolated followed by measuring HDAC activity fluorometrically in a 96-well dark half area opaque microtiter plate. BOC-(Ac)-Lyc-AMC was used as a substrate. **(B)** The cell lysate was isolated from *Cryba1*^fl/fl^ and *Cryba1* cKO mouse RPE cells, followed by a class I HDAC activity assay. Deletion of *Cryba1* in the cKO RPE diminished the enzymatic activity of Class I HDAC. The result was represented as fold change compared with blank value and representative of three individual experiments. (n=3, ***p<0.001). **(C)** Schematic depicting immunoprecipitation of selective HDAC followed by activity assay. HDAC1 **(D)**, HDAC2 **(E)**, HDAC3 **(F),** and HDAC8 **(G)** were immunoprecipitated from *Cryba1*^fl/fl^ and *Cryba1* cKO mouse RPE cells followed by evaluation of activity assay for the respective HDACs. HDAC3 activity was decreased in *Cryba1* cKO compared to *Cryba1*^fl/fl^ **(F)**. The activity of HDAC1/2 and 8 in *Cryba1* cKO was comparable to *Cryba1*^fl/fl^. IgG was used as a negative control. Data has been presented as fold change compared with IgG. Results are representative of three individual experiments. (n=3, ****p<0.0001, NS= not significant). **(H)** Heatmap of enzymatic activity of all class I HDACs (Histone deacetylase) and histone acetylase p300. **(I)** Immunoblot analysis of HDAC1, HDAC2, HDAC3, and HDAC8 from the whole cell lysate isolated from *Cryba1*^fl/fl^ and *Cryba1* cKO mouse RPE cells demonstrate comparable amounts of protein levels in the RPE cells from both genotypes. Actin (cytosolic) and lamin (nuclear) were used as loading controls. The result was representative of three individual experiments. (n=3). **(J)** HDAC3 was immuno-precipitated from *Cryba1*^fl/fl^, βA1 KD and βA3 KO RPE cells, followed by a HDAC3 activity assay. HDAC3 activity significantly decreased in both βA1 KD and βA3 KO cells, while βA1 KD showed a greater reduction in HDAC3 activity than βA3 KO cells. IgG was used as negative control. Data has been presented as fold change compared with IgG value. (n=3, ****p<0.0001 Vs βA1 KD, **p<0.01 Vs βA3 KO). **(K)** Immunoblot analysis of HDAC3 from the total RPE lysate isolated from *Cryba1*^fl/fl^, βA1 KD and βA3 KO. The result was representative of three individual experiments. (n=3). **(L)** NCor1, which is an HDAC3 complex protein, was immuno-precipitated from *Cryba1*^fl/fl^ and *Cryba1* cKO RPE cell lysates followed by HDAC3 activity. The data demonstrates a marked loss of HDAC3’s enzymatic activity in *Cryba1* cKO compared to *Cryba1*^fl/fl^. IgG was used as a negative control. Data has been presented as fold change compared with IgG value. The result was representative of three individual experiments. (n=3, ****p<0.0001). **(M)** SMRT, which is another HDAC3 complex protein, was immuno-precipitated from *Cryba1*^fl/fl^ and *Cryba1* cKO RPE cell lysates followed by HDAC3 activity. The data demonstrates loss of enzymatic activity of HDAC3 in *Cryba1* cKO compared to *Cryba1*^fl/fl^. IgG was used as a negative control. Data has been presented as fold change compared with IgG value. The result was representative of three individual experiments. (n=3, ****p<0.0001). **(N)** Immunoblot analysis of NCor1 and SMRT from the total lysates isolated from *Cryba1*^fl/fl^ and *Cryba1* cKO RPE cells demonstrating comparable protein levels in both lysates, while actin was used as a loading control. The result was representative of three individual experiments. (n=3)

**Figure 2:**
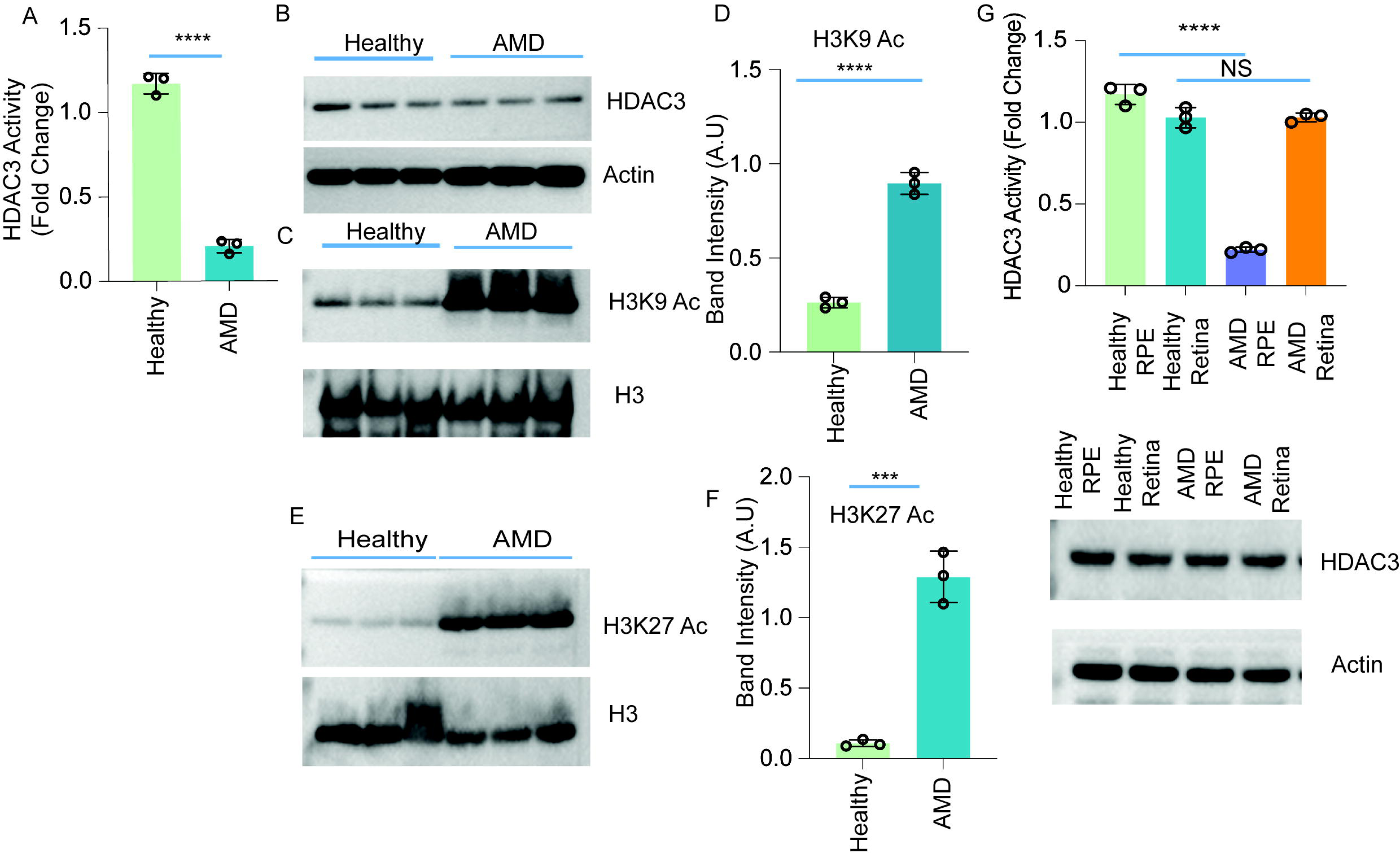
HDAC3 activity is diminished in AMD patients. (A) HDAC3 activity was diminished in AMD patients compared with healthy human individuals. HDAC3 was immunopurified from RPE cells of healthy and AMD patients, followed by a HDAC3 activity assay. Results are representative of three independent experiments. (n=3, ****p<0.0001). **(B)** The HDAC3 protein level in AMD patients was comparable to that of healthy human individuals. Actin was used as a loading control. Results are representative of three independent experiments (n=3). **(C-F)** Western blot analysis of H3K9 and H3K27 acetylation **(C and E)** from healthy and AMD patient RPE cells showing a marked increase in H3k9/27 acetylation in AMD RPE cell lysates when compared with healthy individuals. Respective immunoblots were stripped and then reprobed with an anti-H3 antibody, which was used as a loading control. Results are representative of three independent experiments. **(D and F)** Represent densitometric analysis of H3k9/27 acetylation. n=3 (****p<0.0001, ***p<0.001). **(G)** HDAC3 activity was exclusively diminished in the RPE cells but not in the retina of AMD patients. HDAC3 was immunopurified from RPE cells and retina of healthy patients and those suffering from AMD, followed by a HDAC3 activity assay. The HDAC3 protein level in AMD patients was comparable to healthy individuals. Actin was used as a loading control. Results are representative of three independent experiments. (n=3, ****p<0.0001).

**Figure 3:**
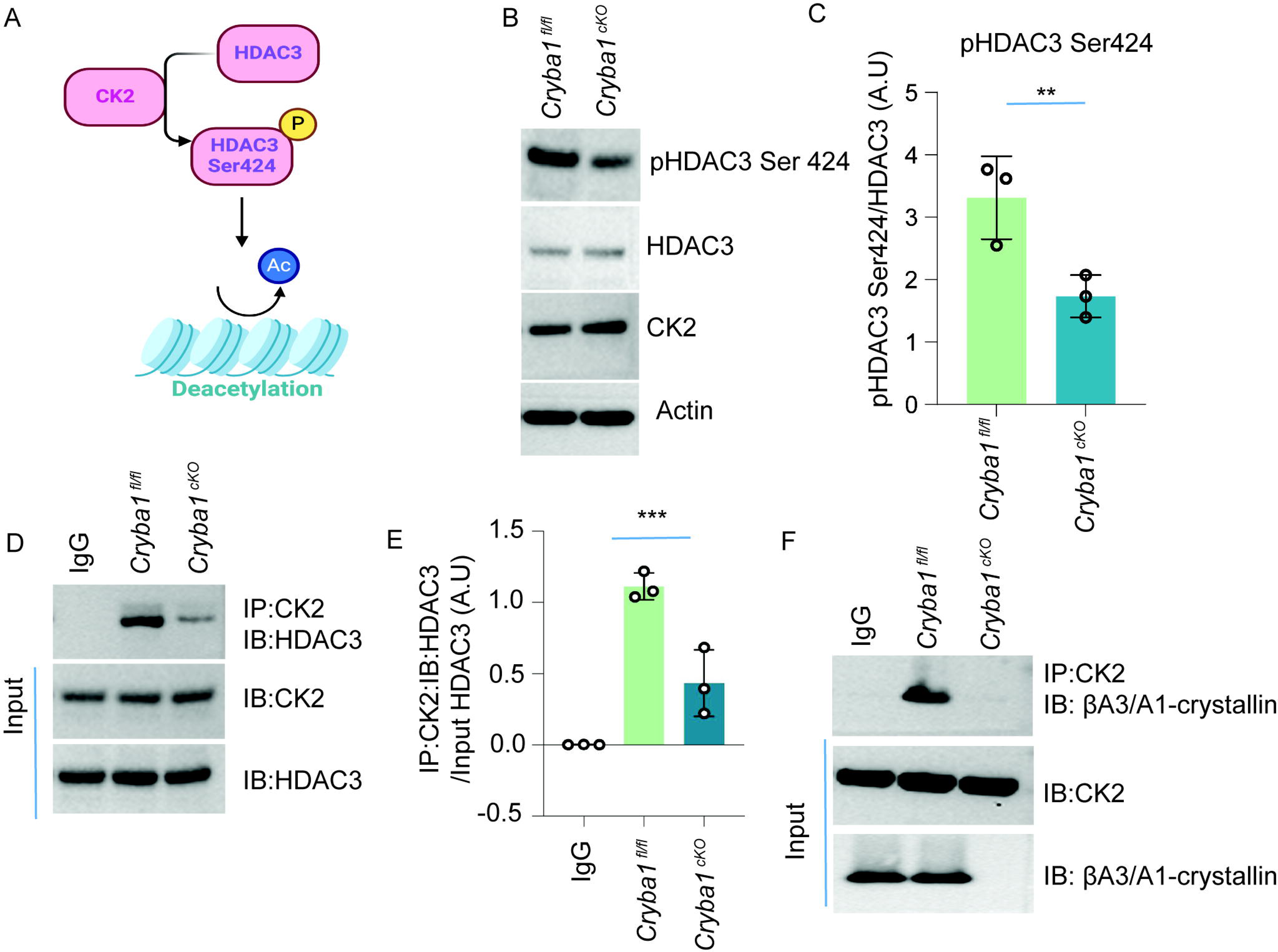
*Cryba1* regulates HDAC3 phosphorylation. (A) Pictorial representation. demonstrates CK2-mediated HDAC3 phosphorylation essential for activating HDAC3’s deacetylase activity. **(B)** Immunoblot analysis shows that *Cryba1* deletion reduced HDAC3 phosphorylation at Serine 424. Actin was used as a loading control. The result was representative of three individual experiments. (n=3) **(C)** Densitometric analysis showing decreased phospho-HDAC3-Ser424 phosphorylation in *Cryba1* cKO compared to *Cryba1*^fl/fl^ (n=3, **p<0.01). **(D)** Endogenous immunoprecipitation study depicted that *Cryba1* deletion diminished endogenous CK2 binding to HDAC3. The result was representative of three individual experiments. (n=3). **(E)** Densitometric analysis confirmed that *Cryba1* deletion diminished endogenous CK2 binding to HDAC3. The result was representative of three individual experiments. (n=3, ***p<0.001) **(F)** Endogenous CK2 binds to *Cryba1* when immunopurified from *Cryba1*^fl/fl^ RPE cells but not *Cryba1* cKO. The result was representative of three individual experiments. (n=3)

HDAC3 requires direct binding to SMRT (silencing mediator of retinoic acid and thyroid hormone receptor) or NCoR1 (nuclear receptor co-repressor 1) corepressor proteins to function as a deacetylase in cells^11^. In our investigation into HDAC3 regulation, we conducted immunoprecipitation of both NCoR1-HDAC3 and SMRT- HDAC3 co-repressor complexes from *Cryba1* cKO and floxed RPE cells. The data demonstrated a substantial reduction (>83.7% and 81.4%) in HDAC3 activity within both complexes in *Cryba1* cKO RPE cells **(Figure 1L, M)** despite no changes in the protein levels of NCoR1 and SMRT **(Figure 1N)**. This suggests that βA3/A1-crystallin likely modulates HDAC3 activity in RPE cells independently of corepressor complexes.

Based on the above findings, our investigation aimed to assess whether individuals suffering from the atrophic form of AMD show any decline in HDAC3 activity with increased histone acetylation. Our data showed that in the RPE of human AMD donor eyes graded by the Minnesota Grading System for disease severity^12^, HDAC3 activity was significantly reduced **(Figure 2A)** relative to age-matched controls, without concurrent changes in HDAC3 protein levels **(Figure 2B)**. In addition, there was a significant increase in the acetylation levels of H3K9 **(Figure 2C, D and Extended Figure 4A, B)** and H3K27 **(Figure 2E, F and Extended Figure 4C, D)** in RPE cells from individuals with AMD. Interestingly, HDAC3 activity was not significantly different in normal and AMD retina samples from the same donors **(Figure 2G).** These findings align with a previous study suggesting that HDAC3-targeted genes in the RPE regulate crucial biological processes associated with AMD^6^. Our study indicates that decreased HDAC3 activity and increased histone acetylation in RPE cells may play a crucial role in AMD, and understanding the underlying mechanism could reveal a novel treatment strategy for AMD.

**Figure 4:**
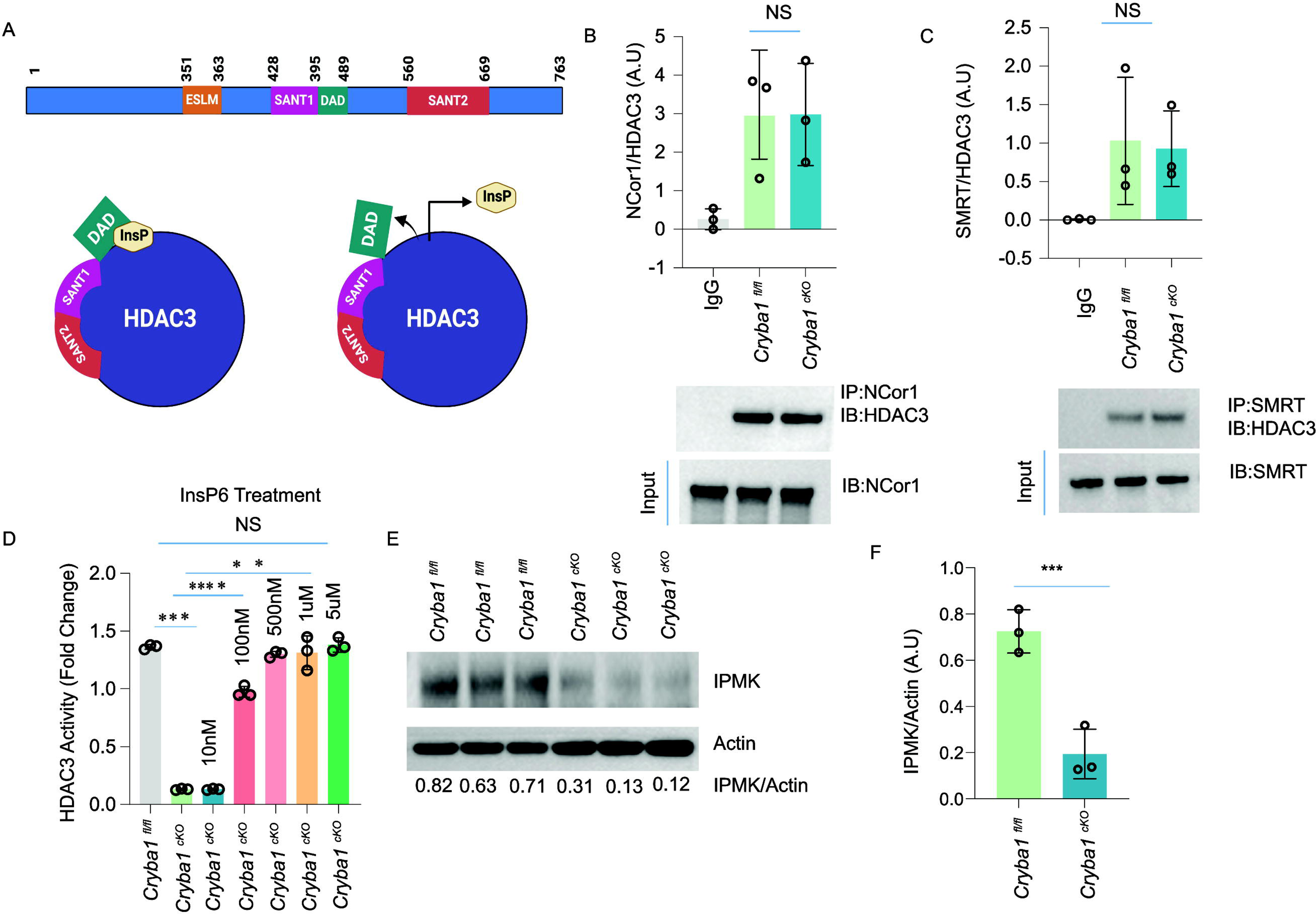
IP6 is essential for HDAC3 activity. (A) Pictorial representation of SMRT. protein domains, which includes the SANT1, SANT2, ESLM (ELM2 specific motif), and DAD domains. The model represents that inositol phosphate (IP) is required for the DAD domain of SMRT protein to interact with HDAC3. **(B)** An endogenous immunoprecipitation study determined that interaction of HDAC3 and Ncor1 in *Cryba1* cKO cells was comparable to *Cryba1*^fl/fl^. The HDAC3/NCor1 complex was pulled down by immunoprecipitating NCor1 followed by western blot for HDAC3. IgG was used as a negative control, and the immunoblot of NCOR1 from the cell lysate was used as an input control. The western blot data is also represented by densitometric analysis. The result was representative of three individual experiments (n=3, NS=not significant). **(C)** The interaction of HDAC3 and SMRT in *Cryba1* cKO cells was comparable to *Cryba1*^fl/fl^. HDAC3 complex integrity with corepressor SMRT was analyzed through immunoprecipitation with *Cryba1*^fl/fl^ and *Cryba1* cKO RPE cell lysates using anti-SMRT antibody followed by immunoblotting for HDAC3. Immunoblotting against anti-SMRT antibodies from respective cell lysates was used as input control, while IgG was used as a negative control. The data is also represented by densitometric analysis. The result was representative of three individual experiments. (n=3, NS=not significant). **(D)** IP6 increased HDAC3 activity in a dose-dependent manner *in vitro* when purified from *Cryba1* cKO cells. To analyze the role of inositol phosphate in HDAC3 activity, endogenous HDAC3 was immunopurified from *Cryba1*^fl/fl^ and *Cryba1* cKO RPE nuclear lysates, followed by dose-dependent IP6 treatment *in vitro*. (n=3, **p<0.01, ***p<0.001, ****p<0.0001). **(E-F)** Immunoblot analysis shows that *Cryba1* deletion diminished IPMK at the protein level in RPE cells from *Cryba1* cKO compared to *Cryba1*^fl/fl^. Actin was used as a loading control. The result was representative of three individual experiments. (n=3, ***p<0.001).

Next, we sought to elucidate the mechanisms through which βA3/A1-crystallin influences the activity of HDAC3. Exploiting the UCSF Chimera program, we observed the potential formation of a complex involving βA3/A1-crystallin (depicted in light blue), HDAC3 (depicted in orange), and the deacetylase-activation-domain (DAD) from the SMRT co-repressor (depicted in green) as illustrated in **(Extended Figure Fig. 5A).** To gain further insight into these protein-protein interactions, co-immunoprecipitation studies in overexpression systems demonstrated a robust binding between βA3/A1- crystallin and HDAC3 **(Extended Figure 5B**). This observation was substantiated by co-purification experiments using endogenous proteins from *Cryba1* floxed RPE cells, indicating a binding interaction, while such interaction was not observed with proteins from *Cryba1* cKO RPE cells **(Extended Figure 5C).** To validate the physical interaction between HDAC3 and βA3/A1-crystallin, an *in vitro* pull-down experiment was conducted. Equal concentrations of recombinant *Cryba1*-myc were incubated with recombinant HDAC3-GST and immunoprecipitated with GST, revealing the binding of both recombinant proteins **(Extended Figure 5D)**. This result confirms the physical interaction between βA3/A1-crystallin and HDAC3, as depicted. To determine the subcellular localization of the *Cryba1*-HDAC3 interaction, we employed a proximity ligation assay (PLA). Confocal microscopy analysis revealed that endogenous βA3/A1- crystallin and HDAC3 proteins interact predominantly within the nucleus. **(Extended Figure 5E)**.

**Figure 5.**
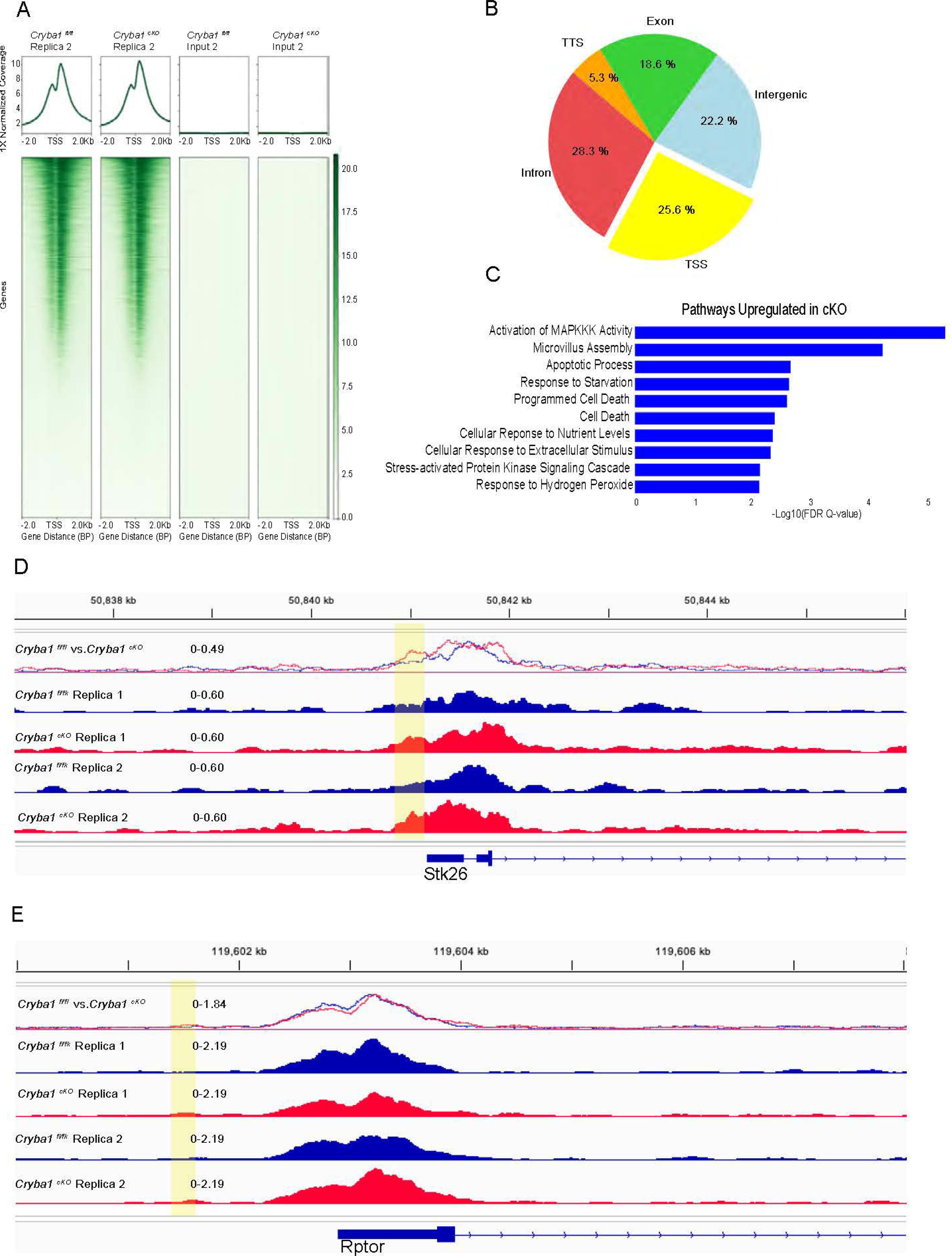
Deletion of *Cryba1* in RPE cells influences epigenetic changes. (A). Heatmaps of chromatin occupancy of the histone marks H3K27Ac in mouse RPE cells. **(B)** Distribution patterns of enriched H3k27ac peaks in *Cryba1* cKO compared to *Cryba1*^fl/fl^. Peaks were filtered using a p-value of <0.05. **(C)** Gene Ontology (GO) enrichment analysis depicting upregulated pathways in *Cryba1* cKO with enriched H3K27 acetylation peaks. **(D)** Genome browser snapshot showing the ChIP seq (acetylated H3K27) track for the Stk26 gene in *Cryba1* cKO. Yellow highlights show the promoter regions. **(E)** Genome browser snapshot showing the ChIP seq (acetylated H3K27) track for the Raptor gene in *Cryba1* cKO. Yellow highlights show the promoter regions.

Next, we wondered how *Cryba1* binding to HDAC3 influences the activation of HDAC3’s deacetylase activity. Previous studies^13^ have proposed that phosphorylation of HDAC3 at serine 424 by Casein kinase 2 (CK2) is essential in activating the enzymatic function of HDAC3 **(Figure 3A)**. Our data showed around a 60% decrease in HDAC3 phosphorylation upon deletion of *Cryba1* in the RPE cells **(Figure 3B-C)**. Remarkably, this reduction was effectively reversed in the *Cryba1* cKO RPE cells by the overexpression of *Cryba1* **(Extended Data** Figure 6A**, B)**. Considering the potential interaction between βA3/A1-crystallin and HDAC3 indicated by protein modeling and pull-down assays, we investigated whether the deletion of *Cryba1* could affect the interaction between CK2 and HDAC3, consequently impacting CK2-mediated HDAC3 phosphorylation. Our data **(Figure 3D, E)** from endogenous pull-down assays demonstrated a marked reduction in the interaction between CK2 and HDAC3 upon *Cryba1* deletion. This suggests that βA3/A1-crystallin may serve as a bridging protein, facilitating interaction between CK2 and HDAC3. To substantiate this premise, we performed additional endogenous pull-down assays confirming the direct interaction between βA3/A1-crystallin and CK2 **(Figure 3F).** In summary, our results suggest that βA3/A1-crystallin may play a crucial role as a linker protein, mediating the interaction between CK2 and HDAC3, thereby influencing HDAC3 phosphorylation and enzymatic activity.

The binding of HDAC3 to its co-repressors, particularly through the DAD domain of the SMRT/NCoR1 protein, is a crucial step that limits HDAC3’s enzymatic activity^14–18^. While the overall interaction between HDAC3 and SMRT/NCoR1 remains intact even in the absence of DAD domain binding, the interaction of HDAC3 with the DAD domain is essential for activating HDAC3’s deacetylase function, which is mediated by highly charged small molecule inositol tetra, penta or hexaphosphate (InsP4/5/6)^14,15,18^ **(Figure 4A)**. Various other domains of the SMRT protein, such as Swi3, Ada2, N-Cor, TFIIIB

(SANT) and ELM-SANT also bind with HDAC3, contributing to the overall interaction of SMRT with HDAC3^15^ **(Figure 4A)**. Intriguingly, despite the loss of HDAC3’s enzymatic activity in *Cryba1* cKO RPE cells, the structural integrity of the HDAC3/SMRT or HDAC3/NCoR1 complexes remains unaffected **(Figure 4B, C)**. It has been reported that higher-order inositol phosphates (HOIP), specifically InsP6 facilitate the interaction between the DAD domain of SMRT and HDAC3^15,16^. The absence of InsP6 binding diminishes HDAC3’s enzymatic activity without affecting the formation of HDAC3/co-regulator complexes. InsP6 functions as an “inter-molecular glue” which binds in a pocket sandwiched between HDAC3 and co-repressor proteins^14,15^. Surprisingly, when we purified HDAC3 from RPE cells lacking βA3/A1-crystallin and incubated it *in vitro,* with varying concentrations of InsP6, we observed a 61.9% rescue of HDAC3 activity at a minimum dose of 100nM **(Figure 4D)** and complete rescue to the wild type (*Cryba1*^fl/fl^) levels at 500 nM **(Figure 4D)**. The data implies that the loss of βA3/A1-crystallin may disrupt HOIP signaling by reducing InsP6 levels, thereby preventing the DAD domain of SMRT or NCoR1 from binding to HDAC3 and ultimately diminishing HDAC3 activity. IPMK (Inositol polyphosphate multi-kinase) is the rate-limiting enzyme in generating HOIPs within mammalian cells and loss of IPMK in cells leads to the substantial reduction of InsP6^19,20^. Notably, the protein expression of IPMK was reduced in *Cryba1* cKO RPE cells **(Figure 4E, F).** This decrease in IPMK expression in *Cryba1* cKO RPE cells suggests an additional mechanism that potentially results in diminished InsP6 levels within these cells. The consequent reduction in InsP6 is assumed to impair HDAC3 activity, thereby affecting various cellular functions regulated by this important enzyme. These findings highlight the intricate relationship between IPMK, InsP6 levels, and HDAC3 activity in the context of *Cryba1* deficiency in the RPE cells.

Prior research has demonstrated the pivotal role of the interaction between HDAC3 and the DAD within NCoR1 and SMRT co-repressors in facilitating the recruitment of HDAC3 to designated gene promoters^17^. This recruitment enables histone deacetylation, a process crucial for gene silencing. Histone deacetylation induces chromatin condensation, rendering it less accessible to transcriptional machinery, thereby suppressing gene transcription^21^. This mechanism is integral for preserving accurate gene expression patterns and sustaining fundamental cellular processes. We conducted ChIP-seq analysis of H3K27ac (as H3k27 acetylation is mostly found to be enriched in gene promoter regions) in *Cryba1*^fl/fl^ and *Cryba1* cKO RPE cells **(Figure 5A and Extended Figure 7A)** to assess the impact of *Cryba1* deletion on the epigenetic landscape. Results showed H3K27 acetylation enrichment at transcription start sites (TSS) for 25.6% of genes in *Cryba1*-deficient cells compared to the floxed RPE cells **(Figure 5B)**. Pathway analysis **(Figure 5C)** of these genes revealed potential regulation of apoptotic, and starvation response pathways, as confirmed by promoter enrichment of H3k27ac in the promoter region of serine/threonine kinase 26 (*Stk26)* **(Figure 5D)** and regulatory-associated protein of mTOR (*Rptor)* **(Figure 5E)**. Raptor, a critical component of mTORC1, plays a pivotal role in regulating cellular metabolism, growth, and autophagy^22^. Recent studies have demonstrated that hyperactivation of mTOR in RPE cells leads to impaired autophagy and elevated oxidative stress, potentially contributing to AMD-like pathology^3, 23,24^. While the STK26 has not been directly implicated in AMD pathogenesis, its involvement in the Hippo signaling pathway—which exhibits crosstalk with mTOR signaling^25,26^—suggests a potential indirect connection. Our findings provide novel insights into these complex molecular interactions, laying the basis for future investigations aimed at elucidating the intricate mechanisms underlying AMD and identifying promising therapeutic targets.

Epigenetic modifications within chromatin play a crucial role in the regulation of various biological processes, and dysregulation of epigenetic mechanisms has been implicated in the development of various complex diseases^27–30^. Specifically, Class I HDACs, such as HDAC3, have been implicated in epigenetic alterations associated with cancer, Alzheimer’s, and Huntington’s diseases^27–29^. However, the involvement of Class I HDACs in the pathogenesis of atrophic AMD has not been previously explored. Our study reveals a significant reduction in HDAC3 activity in individuals with atrophic AMD **(Figure 2A)** and a corresponding mouse model^3–5^ **(Figure 1F)**. While HDAC inhibitors have demonstrated potential in addressing neurodegenerative conditions^28–30^, our research suggests a distinction in the therapeutic approach for AMD compared to Alzheimer’s disease. Specifically, our findings indicate that, in AMD treatment, increasing HDAC3 activation directly in RPE cells may be more efficacious, as we observed no significant alteration in HDAC3 expression but rather a downregulation in its activity. Intriguingly, we found that βA3/A1-crystallin can interact with the active HDAC3 complex in RPE cells, influencing HDAC3 activity. The loss of βA3/A1-crystallin results in decreased HDAC3 activity, a deficit that InsP6 can rescue. InsP6 (phytic acid) is a naturally occurring compound abundant in foods like rice, potatoes, and legumes and has been implicated in various cellular processes, including chromatin remodeling and gene expression^31–33^. Here, we show that InsP6 may play a role in activating HDAC3 activity in RPE cells **(Figure 4D-F)** by modulating the complex formation of HDAC3 with DAD and βA3/A1-crystallin. Furthermore, we discovered that CK2- mediated HDAC3 phosphorylation, which is crucial for HDAC3’s enzymatic activity, was impaired in *Cryba1*-deficient cells **(Figure 3A-F)**. We identified that βA3/A1-crystallin acts as a linker protein, and the loss of *Cryba1* diminished the HDAC3-CK2 interaction **(Figure 3D-F)**. Notably, previous research has established that InsP6 directly interacts with CK2, enhancing its kinase activity. This observation, coupled with our results, suggests a plausible mechanism wherein reduced InsP6 levels in *Cryba1*-deficient cells may impair HDAC3 activation through two concurrent pathways: (1) disruption of DAD domain binding to HDAC3 and (2) partial inhibition of CK2 kinase activity.

This novel regulation of HDAC3 in RPE cells by βA3/A1-crystallin unveils an unexplored function of the crystallin in RPE cells, which may underlie its established role in lysosomal function, mTOR signaling, and inflammation^3–5^. Importantly, our results suggest that elevating βA3/A1-crystallin levels or InsP6 treatment in diseased RPE cells may mitigate multiple dysregulated pathways by influencing epigenetic changes associated with AMD, a multi-factorial disease that currently has limited therapeutic options^1–2^.

## Methods Animals

All animal studies were conducted in accordance with the Guide for the Care and Use of Animals (National Academy Press) and were approved by the University of Pittsburgh Animal Care and Use Committee. Both male and female mice were used in this study. RPE-specific βA3/A1-crystallin conditional (*Cryba1* cKO), as well as global βA3 KO and βA1 KD mice were generated on a C57BL/6J background as previously described^3–5,10^. All mice that were used in this study were negative for RD8 mutation^3^. Animals were euthanized as per approved guidelines and the RPE was harvested for downstream experiments (immunoblot, HDAC activity assays) as explained previously^3–5^.

### RPE explant culture and Pan-HDAC Inhibitor SAHA treatment as well as adenovirus-*Cryba1* construct infection

Eyes from 4-5 month-old *Cryba1* floxed and cKO mice were enucleated, and the RPE-choroid-sclera (RCS) complex was harvested and placed onto PVDF membranes after flattening them by making several relaxing cuts^5,34^. The explants were then cultured face up in complete media as previously described^5,34^. *Cryba1*^fl/fl^ RPE explants were incubated with 2 mM of SAHA (Sigma Aldrich, Cat# SML0061-5MG) and incubated overnight followed by estimation of histone acetylation by immunoblot^3–5,19^. *Cryba1* cKO RPE explants were also infected with either adenovirus(Ad)-*Cryba1* construct or vehicle control at a dose of 10^5^ vg/ml for 24 h and subsequent experiments were performed.

### Human AMD donors

The human AMD donor sections were gifts from Dr. Flores-Bellver’s laboratory at the University of Colorado. The donors were all Caucasian decent and had an average age of 78 ± 3 years. The control donors did not have any eye diseases. Both sexes were included for both control (n=6) and AMD (n=6) donors. The disease was staged according to the Minnesota Grading System (MGS), and all the AMD donors were classified as MGS2^12^.

### Western blot analysis

Cell lysates were prepared through syringe flush by using a salt-free (NaCl-free) Lysis buffer (50 mM Tris pH 7.5, 50 mM potassium acetate, 5 % v/v glycerol, 0.3 % v/v Triton X-100, one tablet of Roche complete protease inhibitor). We used salt-free (NaCl-free) lysis buffer for all the assays because high a salt concentration dissociates higher-order inositol phosphate (HOIPs) from HDAC3/1, which is essential for their deacetylase activity^15,16^. Samples were centrifuged at 15,000 g for 10 min, and the protein concentration of the supernatant was measured. Proteins were resolved by SDS-polyacrylamide gel electrophoresis (NuPAGE Bis-Tris Midi GEL, Life Technologies, Cat #: WG1402BX10) and transferred to Immobilion-P PVDF (Millipore-Sigma, Cat #: IPVH00010) transfer membranes. The membranes were incubated with primary antibody diluted in 3% BSA in Tris- Buffered Saline with Tween 20 (20 mM Tris-HCl, pH 7.4, 150 mM NaCl, and 0.02% Tween 20) overnight incubation at 4°C. Respective antibodies for western blot anti-Cryba1 (Abcam, Cat #: ab151722), anti-HDAC3 (Santa Cruz Biotechnology, Cat #: sc-376957), anti-HDAC1 (Santa Cruz Biotechnology, Cat #: sc-81598), anti-HDAC2 (Santa Cruz Biotechnology, Cat #: SC-7899), anti-HDAC8 (Biolegend, Cat #: 685504), anti-pHDAC3 Ser424 (Thermo Fischer Scientific, Cat #: PA5-99339), anti-CK2 (ProteinTech, Cat #: 10992-1-AP), anti-beta actin (ProteinTech, Cat #: 81115-1-RR), anti-lamin (ProteinTech, Cat #: 12987-1-AP), anti-SMRT (Bethyl Laboratories, Cat #: A301-148A), anti-Ncor1 (Cell Signaling Technology, Cat #: 5948), anti-H3 (Cell Signaling Technology, Cat #: 12648), anti-H3k8Ac (Cell Signaling Technology, Cat #: 2598), anti-H3k9Ac (Cell Signaling Technology, Cat #: 9649), anti- H3k18Ac (Cell Signaling Technology, Cat #: 13998), anti-H3k27Ac (Abcam, Cat #: ab4729) ,anti-H3k56Ac (Cell Signaling Technology, Cat #: 4243), anti-H4k8Ac (Cell Signaling Technology, Cat #: 2594), anti-H4k12Ac (Cell Signaling Technology, Cat #: 13944), anti-H4k16Ac (EMD Millipore Corp., Cat #: 07-329) and anti-Myc (ProteinTech, Cat #:16286-1-AP). Following primary antibody incubation, the PVDF membrane was washed three times with Tris-buffered saline/Tween-20, and incubated with HRP-conjugated secondary antibody (ECL, Cat #s: NA934V and NA931V), and the bands visualized by chemiluminescence (Super Signal West Pico, Pierce, Cat #: 34579)^4,19^. The depicted blots are representative replicates selected from at least three experiments. Densitometric analysis was performed using ImageJ software. Blots for acetylated histones were stripped using Restore Western Blot stripping buffer (Thermo Scientific, Cat # 21059), then reprobed with H3 antibody (Cell Signaling Technology, Cat #: 12648S) at room temperature for 2 h followed by secondary antibody incubation and development.

### HAT Activity

HAT activity was measured by following the HAT Assay Kit (Abcam, Cat #: ab-204709) protocol. *Cryba1*^fl/fl^ and *Cryba1* cKO cells were lysed in salt-free (NaCl-free) lysis buffer (50 mM Tris pH 7.5, 50 mM potassium acetate, 5 % v/v glycerol, 0.3 % v/v Triton X-100, Roche complete protease inhibitor). Samples were centrifuged at 15,000 g for 10 min, and the protein concentration of the supernatant was measured. About 20 μg total protein from the lysate was incubated against anti-p300 CBP HAT antibody (Cell Signaling Technology, Cat #: 86377) respectively at 4°C for 1 hr, followed by capturing the antibody with EZview A/G beads (Millipore-Sigma, Cat #: E3403). Beads were washed three times with an ice-cold reaction buffer (supplied with kit). Immunoprecipitated P300 was used as a protein source. Absorbance was measured after specific substrate reaction by using a Spectramax iD5 plate reader (Molecular Devices). All measurements were performed in triplicate and data analyzed using GraphPad Prism (version 6.0, GraphPad Software, Inc).

### HDAC activity

To study global HDAC activity from *Cryba1*^fl/fl^ and *Cryba1* cKO cell lysates, we used an HDAC activity assay Kit (Active Motif, Cat #: 56200). In brief, we lysed cells in a low-salt lysis buffer, as stated before, followed by centrifugation at 15,000 g to isolate the supernatant. About 2 μg of protein from *Cryba1*^fl/fl^ and *Cryba1* cKO lysates were incubated with the substrate provided with the kit, and then fluorescence was measured using a plate reader.

To study individual HDAC activity, the HDAC1, 2, 3 and 8 were individually immunopurified, followed by an activity assay using the following kits: HDAC Assay Kit (Active Motif, Cat#: 56200 for HDAC1, 2 and 8) and the HDAC3 Assay Kit (BPS, Cat#: 10186-628). In brief, *Cryba1*^fl/fl^ and *Cryba1* cKO cells were lysed in salt-free (NaCl-free) lysis buffer (50 mM Tris pH 7.5, 50 mM potassium acetate, 5 % v/v glycerol, 0.3 % v/v Triton X-100, one tablet of Roche complete protease inhibitor). Then, the lysate was centrifuged at 15,000 g for 10 min, and the protein concentration of the supernatant was measured. Around 20 μg total protein from the lysate was incubated against anti- HDAC1 (Santa Cruz Biotechnology, Cat # sc-81598), anti-HDAC2 (Santa Cruz Biotechnology, Cat #: sc-7899), anti-HDAC3 (Santa Cruz Biotechnology, Cat #: sc- 81600) and anti-HDAC8 (Biolegend, Cat #: 685504) antibodies, respectively, at 4°C for 1 hr followed by capturing antibody with EZview A/G beads (Millipore Sigma, Cat #: E3403). Beads were washed 3x with an ice-cold reaction buffer (supplied with the kit). After a specific substrate reaction, fluorescence was measured using a Spectra max iD5 plate reader (Molecular Devices). All measurements were performed in triplicate, and data was analyzed using GraphPad Prism (version 6.0, GraphPad Software, Inc)^14,16^.

### InsP6 Rescuing HDAC3 activity

*Cryba1* cKO cells were lysed in salt-free (NaCl-free) lysis buffer (50 mM Tris pH 7.5, 50 mM potassium acetate, 5 % v/v glycerol, 0.3 % v/v Triton X-100, and one Roche complete protease inhibitor tablet) because high salt concentration dissociates higher-order inositol phosphate (HOIPs) from HDAC3/1, which is essential for their deacetylase activity^15,16^. Samples were centrifuged at 15,000 g for 10 min, and the protein concentration of the supernatant was measured. 20 μg total protein from the lysate was incubated against anti-HDAC3 antibodies respectively at 4°C for 1 hr, followed by capturing antibodies with EZview A/G beads (Millipore Sigma, Cat #: E3403). Beads were washed 3x with an ice-cold reaction buffer (supplied with the kit). Following washing, beads were resuspended in reaction buffer and incubated for 1 hr at room temperature with 10nM, 100nM, 500nM, and 1μM concentrations of IP6 (Sigma Aldrich Cat #: P8810-100G). After incubation, beads were washed 3X with reaction buffer. HDAC3 activity was measured by the addition of a fluorogenic substrate provided by the HDAC3 Assay Kit (BPS, Cat #: 10186-628) and read using a Spectramax iD5 plate reader (Molecular Devices). All measurements were performed in triplicate data analyzed, and the IC50 value was calculated using GraphPad Prism (version 6.0, GraphPad Software, Inc)^14,16^.

### Immunoprecipitation

To ascertain HDAC3 and βA3/A1-crystallin binding in overexpression conditions, *Cryba1*^fl/fl^ RPE explants were infected with Ad-RFP (Vector Biolabs, Cat # 1660) or Ad- *Cryba1*-RFP (Vector Biolabs, customized) constructs at 10^5^ vg/ml for 24 h^5^. The RPE lysates were prepared^3–5^, and immunoprecipitation was performed using anti-RFP beads, followed by immunoblot for HDAC3. The whole cell lysates were also used to immunoblot for HDAC3 and RFP as input controls. To analyze endogenous binding, RPE cells were lysed in salt-free lysis buffer (50 mM Tris pH 7.5, 50 mM potassium acetate, 5 % v/v glycerol, 0.3 % v/v Triton X-100, and one Roche complete protease inhibitor tablet). For the endogenous immunoprecipitation (IP) study, respective antibodies against βA3/A1-crystallin, CK2, or HDAC3 were used, followed by western blot of the binding partners. In brief, the IP was performed from 500 μg of protein lysate. Protein lysates were incubated for 2 hr at 4°C with respective antibodies, then the antibody-protein complex was captured with EZview A/G beads (Millipore Sigma, Cat #: E3403). Beads were pelleted and washed with lysis buffer 3x, followed by elution in sample buffer. Immunoprecipitated samples were resolved on a NuPAGE Bis-Tris gel, followed by western blotting.

### *In vitro* binding assay

Equal amount of recombinant myc-βA3/A1-crystallin (Origene, Cat #: TP321965) was co-incubated with either recombinant GST-HDAC3 (Signal Chem, Cat #: H85-30G) or GST (Signal Chem, Cat #: G52-30U) in lysis buffer and the complex was maintained for 30 min at 4°C. After the addition of GST beads, incubation continued for an additional 45 min and washed 3x with ice cold lysis buffer. SDS sample buffer was added and binding was confirmed by western blotting of anti-Myc antibodies^19^.

### Proximity ligation assay (PLA)

Proximity ligation assay was carried out on RPE flat mounts following the manufacturers’ protocol (Navinci Diagnostics, NaveniFlex Tissue MR Red, Cat #: NT.MR.100). Antibody for HDAC3 (Cell Signaling Technology, Cat #: 3949,) was used at a 1:100 dilution with beta crystallin A3 (Abcam, Cat #: ab-151722), for the first set of PLA to check the interaction between crystallin and HDAC3.

### Molecular modelling

Models of molecular structure for human βA3/A1-crystallin, HDAC3 and SMRT-DAD were obtained from the AlphaFold Protein Structure Database (https://alphafold.ebi.ac.uk/). Molecular modeling of the possible interaction between these proteins was done using the ‘movement’ tool in the extensible molecular modeling system UCSF CHIMERA, VER. 1.17.1 (https://www.cgl.ucsf.edu/chimera/)

### RNA-seq data analysis

The following published RNA-Seq datasets from 5-month-old mouse RPE cells were downloaded^35^: GSM4043956 (*Cryba1*^fl/fl^ (WT)-5 Months) and GSM4043957 (*Cryba1* cKO-5 Months). The FASTQ files were analyzed in Partek (v 12.0.0)^36^. All pre- and post-alignment QA/QC was performed in Partek. Reads were aligned to whole mm10 genome using STAR (v 2.7.8a). Aligned reads were quantified to mm10 Ensembl Transcripts release 102 using HTSeq (v 0.11.0). Differential gene expression was performed using DESeq2 (v 3.5).

### Chromatin Immunoprecipitation (ChIP)

Dissected RPE were incubated with 1% formaldehyde in PBS at room temperature for 10 min. Fixation was stopped by the addition of 1X glycine. TruChIP Chromatin Shearing Kit (Covaris, Cat #: 520127) was used to shear chromatin. Chromatin immunoprecipitation (ChIP) was performed using iDeal ChIPseq kit (Diagenode, Cat #: C01010051) on 25 μg of sheared chromatin using 5 μl of H3K27-Ac antibody (Abcam, Cat #: ab-4729). Crosslinking was reversed by overnight incubation at 65°C using proteinase K treatment. DNA was then purified using MinElute Reaction Cleanup Kit (Qiagen, Cat #: 28204). To confirm the significant enrichment, qPCR was performed on control genes. Libraries of samples were prepared using a NEBNext Ultra™ II DNA Library Prep Kit (NEB, Cat #: E7645). The library was sequenced using single-end reads (150 base pair reads) on a NextSeq 500 system at the ULNV/NIPM Genomics Core.

### ChIP-seq data analysis

FASTQ files from sequencing were aligned to the Mouse mm10 genome in Partek (v 12.0.0)^36^ using BWA-MEM (v 0.7.17). The peaks were identified using MACS2 (v 3.0.0a7), broad region and q-value cutoff of 0.05, compared with respective input samples and annotated using mm10 Ensembl Transcripts release 102. Differential peak analysis was performed comparing cKO H3K27 vs. FL H3K27 samples using DESeq2 (v 3.5). Peaks with a p-value <= 0.05 were considered significant.

To generate TSS plots, BAM files from alignment performed in Partek were downloaded and converted to bigwig files with DeepTools (v 3.5.5)^37^ bamCoverage function. Reads were extended to a fragment length of 200 base pairs. Read coverage was calculated using a 10-base pair window and normalized to 1x depth using an effective genome size of 2,650,000,000. Chromosome X was ignored for normalization and blacklisted regions were downloaded from https://github.com/Boyle-Lab/Blacklist^38^

and excluded. Count matrices for all samples were generated with the DeepTools compute Matrix function using a bin size of 10. TSS were downloaded from the UCSC Table Browser for mm10 NCBI RefSeq Track (https://genome.ucsc.edu/cgi-bin/hgTables)^39^. The reference point was 2000 base pairs above and below TSS. DeepTools plot Heatmap function was used to generate TSS plots.

To generate chromosome plots, FL H3K27 samples were averaged with DeepTools bigwigCompare function. The same operation was performed for *Cryba1* cKO H3K27 samples. Integrative Genomics Viewer (v 2.17.4) was used to generate choromsome plot views (https://igv.org/)^40^.

### Pathway analysis

Pathway analysis was performed on significant peaks (p-value <= 0.05) with GREAT v4.0.4^41^ using mouse GRCm38 assembly (http://great.stanford.edu/public/html/). The whole genome was used as the background region. The whole genome was used as the reference set.

### Statistical analysis

All plots and statistical analyses were performed with Prism 9 (GraphPad) software^3–5^. Statistical significance was determined by either Student’s t-test (two-tailed) for two groups or 1-way ANOVA for multiple groups with similar samples^3–5,19^. Error bars represent the standard deviation of the mean and indicate replicates or the number of animals employed^3–5,19^. Results were representative of at least three independent experiments (n). Differences between groups were considered significant when *p<0.01, **p<0.001, and ***p<0.0001.

## Acknowledgements

We would like to thank J. Samuel Zigler, Jr. for his help during the preparation of the manuscript and the NIPM genomic core facility for the NGS studies. This work was supported by NIH 5P20GM121325 COBRE grant and University of Nevada, Las Vegas start-up funds to Prasun Guha. This study was also supported by NIH 1R01EY031594- 01A1 (DS), NIH 1R01EY032516-01A1 (DS), University of Pittsburgh start-up funds (DS), Edward N. & Della L. Thome Memorial Foundation Awards Program in Age-Related Macular Degeneration (DS), NIH K99EY033421 (SG), P30 core award EY08098 from the National Eye Institute, NIH (to the University of Pittsburgh Department of Ophthalmology), and unrestricted funds from The Research to Prevent Blindness Inc., NY (to the University of Pittsburgh Department of Ophthalmology). DS was the Jennifer Salvitti Davis, M.D. Professor at the University of Pittsburgh.

## Author contributions

Prasun Guha and DS designed the study. SC performed most histone acetylation related experiments and biochemical assays. ZS, and LVP performed NGS data analysis. LVP and NT performed biochemical experiments. SG, SB, and Pooja Gautam, generated, genotyped, and isolated RPE cells for the *Cryba1*^fl/fl^, *Cryba1* cKO, βA1 KD and βA3 KO mice. All assays conducted in this research utilized these cells. SG conducted the binding assay, while SB performed the immunofluorescence assay. Additionally, SG and SB did the RPE explant cultures for *in vitro* experiments, both with and without specific treatments. ED and IAD did the ChIP assay. YS generated the computer model. MFB provided the human tissues and characterized them. SC and SH designed the figures. DS and Prasun Guha wrote the manuscript. All authors reviewed the results and approved the final version of the manuscript.

## References

1. Boya P, Kaarniranta K, Handa JT, et al. Lysosomes in retinal health and disease.

2. *Trends* *Neurosci*. 2023;46(12):1067#1082. doi: 10.1016/j.tins.2023.09.006.

2. Handa JT, Bowes Rickman C, Dick AD, et al. A systems biology approach towards understanding and treating non-neovascular age-related macular degeneration. Nat Commun. 2019;10(1):3347. doi: 10.1038/s41467-019-11262-1.

3. Valapala M, Wilson C, Hose S, et al. Lysosomal-mediated waste clearance in retinal pigment epithelial cells is regulated by CRYBA1/betaA3/A1-crystallin via V-ATPase- MTORC1 signaling. Autophagy. 2014;10(3):480–496. doi: 10.4161/auto.27292.

4. Ghosh S, Shang P, Terasaki H, et al. A role for betaA3/A1-crystallin in type 2 EMT of RPE cells occurring in dry age-related macular degeneration. Invest Ophthalmol Vis Sci. 2018;59(4):AMD104-AMD113. doi: 10.1167/iovs.18-24132.

5. Shang P, Stepicheva N, Teel K, et al. βA3/A1-crystallin regulates apical polarity and EGFR endocytosis in retinal pigmented epithelial cells. Commun Biol. 2021;4(1):850. doi: 10.1038/s42003-021-02386-6.6.

6. Wang J, Zibetti C, Shang P, et al. ATAC-seq analysis reveals a widespread decrease of chromatin accessibility in age-related macular degeneration. Nat Commun. 2018;9(1):1364-y. doi: 10.1038/s41467-018-03856-y.

7. Rodems TS, Heninger E, Stahlfeld CN, et al. Reversible epigenetic alterations regulate class I HLA loss in prostate cancer. Commun Biol. 2022;5(1):897–6. doi: 10.1038/s42003-022-03843-6.

8. Shi X, Ding W, Li T, Zhang Y, Zhao S. Histone deacetylase (HDAC) inhibitor, suberoylanilide hydroxamic acid (SAHA), induces apoptosis in prostate cancer cell lines via the akt/FOXO3a signaling pathway. Med Sci Monit. 2017;23:5793–5802. doi: 10.12659/msm.904597.

9. Dancy BM, Cole PA. Protein lysine acetylation by p300/CBP. Chem Rev. 2015;115(6):2419–2452. doi: 10.1021/cr500452k.

10. Ghosh S, Liu H, Yazdankhah M, et al. betaA1-crystallin regulates glucose metabolism and mitochondrial function in mouse retinal astrocytes by modulating PTP1B activity. Commun Biol. 2021;4(1):248–5. doi: 10.1038/s42003-021-01763-5.

11. Mottis A, Mouchiroud L, Auwerx J. Emerging roles of the corepressors NCoR1 and SMRT in homeostasis. Genes Dev. 2013;27(8):819–835. doi: 10.1101/gad.214023.113.

12. Olsen TW, Liao A, Robinson HS, Palejwala NV, Sprehe N. The nine-step minnesota grading system for eyebank eyes with age related macular degeneration: A systematic approach to study disease stages. Invest Ophthalmol Vis Sci. 2017;58(12):5497–5506. doi: 10.1167/iovs.17-22161.

13. Zhang X, Ozawa Y, Lee H, et al. Histone deacetylase 3 (HDAC3) activity is regulated by interaction with protein serine/threonine phosphatase 4. Genes Dev. 2005;19(7):827–839. doi: 10.1101/gad.1286005.

14. Watson PJ, Millard CJ, Riley AM, et al. Insights into the activation mechanism of class I HDAC complexes by inositol phosphates. Nat Commun. 2016;7:11262. doi: 10.1038/ncomms11262.

15. Millard CJ, Watson PJ, Celardo I, et al. Class I HDACs share a common mechanism of regulation by inositol phosphates. Mol Cell. 2013;51(1):57–67. doi: 10.1016/j.molcel.2013.05.020.

16. Watson PJ, Fairall L, Santos GM, Schwabe JWR. Structure of HDAC3 bound to co- repressor and inositol tetraphosphate. Nature. 2012;481(7381):335-340. doi: 10.1038/nature10728.

17. You S, Lim H, Sun Z, Broache M, Won K, Lazar MA. Nuclear receptor co-repressors are required for the histone-deacetylase activity of HDAC3 in vivo. Nat Struct Mol Biol. 2013;20(2):182–187. doi: 10.1038/nsmb.2476.

18. Guenther MG, Barak O, Lazar MA. The SMRT and N-CoR corepressors are activating cofactors for histone deacetylase 3. Mol Cell Biol. 2001;21(18):6091–6101. doi: 10.1128/MCB.21.18.6091-6101.2001.

19. Guha P, Tyagi R, Chowdhury S, et al. IPMK mediates activation of ULK signaling and transcriptional regulation of autophagy linked to liver inflammation and regeneration. Cell Rep. 2019;26(10):2692–2703.e7. doi: 10.1016/j.celrep.2019.02.013.

20. Kim E, Beon J, Lee S, et al. IPMK: A versatile regulator of nuclear signaling events. Adv Biol Regul. 2016;61:25–32. doi: 10.1016/j.jbior.2015.11.005.

21. Watts BR, Wittmann S, Wery M, et al. Histone deacetylation promotes transcriptional silencing at facultative heterochromatin. Nucleic Acids Res. 2018;46(11):5426–5440. doi: 10.1093/nar/gky232.

22. Saxton RA, Sabatini DM. mTOR Signaling in Growth, Metabolism, and Disease.

24. Cell. 2017;168(6):960#976. doi: 10.1016/j.cell.2017.02.004.

23. Cai J, Litwin C, Cheng R, et al. DARPP32, a target of hyperactive mTORC1 in the retinal pigment epithelium. Proc Natl Acad Sci U S A. 2022;119(33):e2207489119. doi: 10.1073/pnas.2207489119.

24. Huang J, Gu S, Chen M, et al. Abnormal mTORC1 signaling leads to retinal pigment epithelium degeneration. Theranostics. 2019;9(4):1170–1180. doi: 10.7150/thno.26281.

27. Wang D, He J, Huang B, et al. Emerging role of the Hippo pathway in autophagy.

28. *Cell Death* *Dis*. 2020;11(10):880. doi: 10.1038/s41419-020-03069-6.

26. Honda D, Okumura M, Chihara T. Crosstalk between the mTOR and Hippo pathways. Dev Growth Differ. 2023;65(6):337–347. doi: 10.1111/dgd.12867.

27. Rodems TS, Heninger E, Stahlfeld CN, et al. Reversible epigenetic alterations regulate class I HLA loss in prostate cancer. Commun Biol. 2022;5(1):897–6. doi: 10.1038/s42003-022-03843-6.

28. Janczura KJ, Volmar C, Sartor GC, et al. Inhibition of HDAC3 reverses alzheimer’s disease-related pathologies in vitro and in the 3xTg-AD mouse model. Proc Natl Acad Sci U S A. 2018;115(47):E11148–E11157. doi: 10.1073/pnas.1805436115.

29. Jia H, Morris CD, Williams RM, Loring JF, Thomas EA. HDAC inhibition imparts beneficial transgenerational effects in huntington’s disease mice via altered DNA and histone methylation. Proc Natl Acad Sci U S A. 2015;112(1):56. doi: 10.1073/pnas.1415195112.

30. Chuang D, Leng Y, Marinova Z, Kim H, Chiu C. Multiple roles of HDAC inhibition in neurodegenerative conditions. Trends Neurosci. 2009;32(11):591–601. doi: 10.1016/j.tins.2009.06.002.

31. Pujol A, Sanchis P, Grases F, et al. Phytate Intake, Health and Disease: "Let Thy Food Be Thy Medicine and Medicine Be Thy Food". Antioxidants (Basel). 2023;12(1):146. doi: 10.3390/antiox12010146.

32. Beon J, Han S, Yang H, et al. Inositol polyphosphate multikinase physically binds to the SWI/SNF complex and modulates BRG1 occupancy in mouse embryonic stem cells. Elife. 2022;11:10.7554/eLife.73523. doi: 10.7554/eLife.73523.

33. Reilly L, Semenza ER, Koshkaryan G, et al. Loss of PI3k activity of inositol polyphosphate multikinase impairs PDK1-mediated AKT activation, cell migration, and intestinal homeostasis. iScience. 2023;26(5):106623. doi: 10.1016/j.isci.2023.106623.

34. Shang P, Stepicheva NA, Liu H, et al. A Novel Method of Mouse RPE Explant Culture and Effective Introduction of Transgenes Using Adenoviral Transduction for In Vitro Studies in AMD. Int J Mol Sci. 2021;22(21):11979. doi: 10.3390/ijms222111979.

35. Ghosh S, Padmanabhan A, Vaidya T, et al. Neutrophils homing into the retina trigger pathology in early age-related macular degeneration. Commun Biol. 2019;2:348. doi: 10.1038/s42003-019-0588-y.

36. Fonseca TL, Garcia T, Fernandes GW, et al. Neonatal thyroxine activation modifies epigenetic programming of the liver. Nat Commun. 2021;12;4446. 10.1038/s41467-021-24748-8.

37. Ramírez F, Ryan DP, Grüning B, Bhardwaj V, Kilpert F, Richter AS, Heyne S, Dündar F, Manke T. deepTools2: a next generation web server for deep-sequencing data analysis. Nucleic Acids Res. 2016;44(W1):W160–5. doi: 10.1093/nar/gkw257.

38. Amemiya HM, Kundaje A, Boyle AP. The ENCODE Blacklist: Identification of Problematic Regions of the Genome. Sci Rep. 2019;9(1):9354. doi: 10.1038/s41598-019-45839-z.

39. Nassar LR, Barber GP, Benet-Pagès A, et al. The UCSC Genome Browser database: 2023 update. Nucleic Acids Res. 2023;51(D1):D1188–D1195. doi: 10.1093/nar/gkac1072.

40. Robinson JT, Thorvaldsdóttir H, Winckler W, et al. Integrative genomics viewer. Nat Biotechnol. 2011;29(1):24#6. doi: 10.1038/nbt.1754.

41. McLean CY, Bristor D, Hiller M, et al. GREAT improves functional interpretation of cis-regulatory regions. Nat Biotechnol. 2010;28(5):495–501. doi: 10.1038/nbt.1630.

