## Extended Data for "βA3/A1-crystallin is an epigenetic regulator of histone deacetylase 3 (HDAC3) in the retinal pigmented epithelial (RPE) cells"

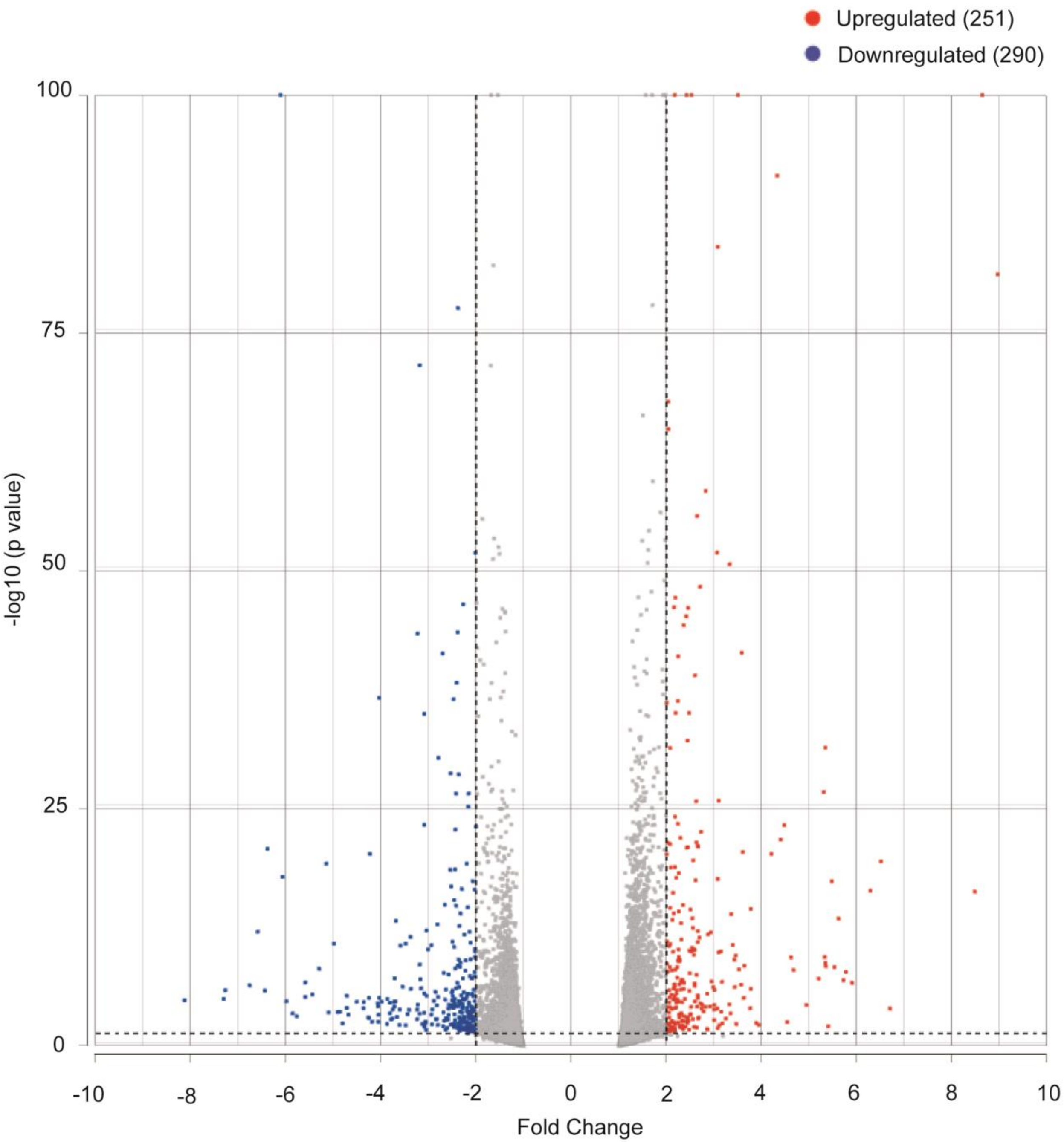


**<u>Extended Figure 1</u>: *Cryba1* deletion impacts the global transcriptome in mouse RPE cells.** Volcano plot (log 10[*p*-value] vs. Fold change) displays differentially expressed genes after *Cryba1* deletion. Red dots represent upregulated gene expression, while blue dots represent downregulated gene expression. The y-axis denotes –log10 p Value while the x-axis denotes log2 fold change value. The result is representative of eight individual experiments. (n=8)

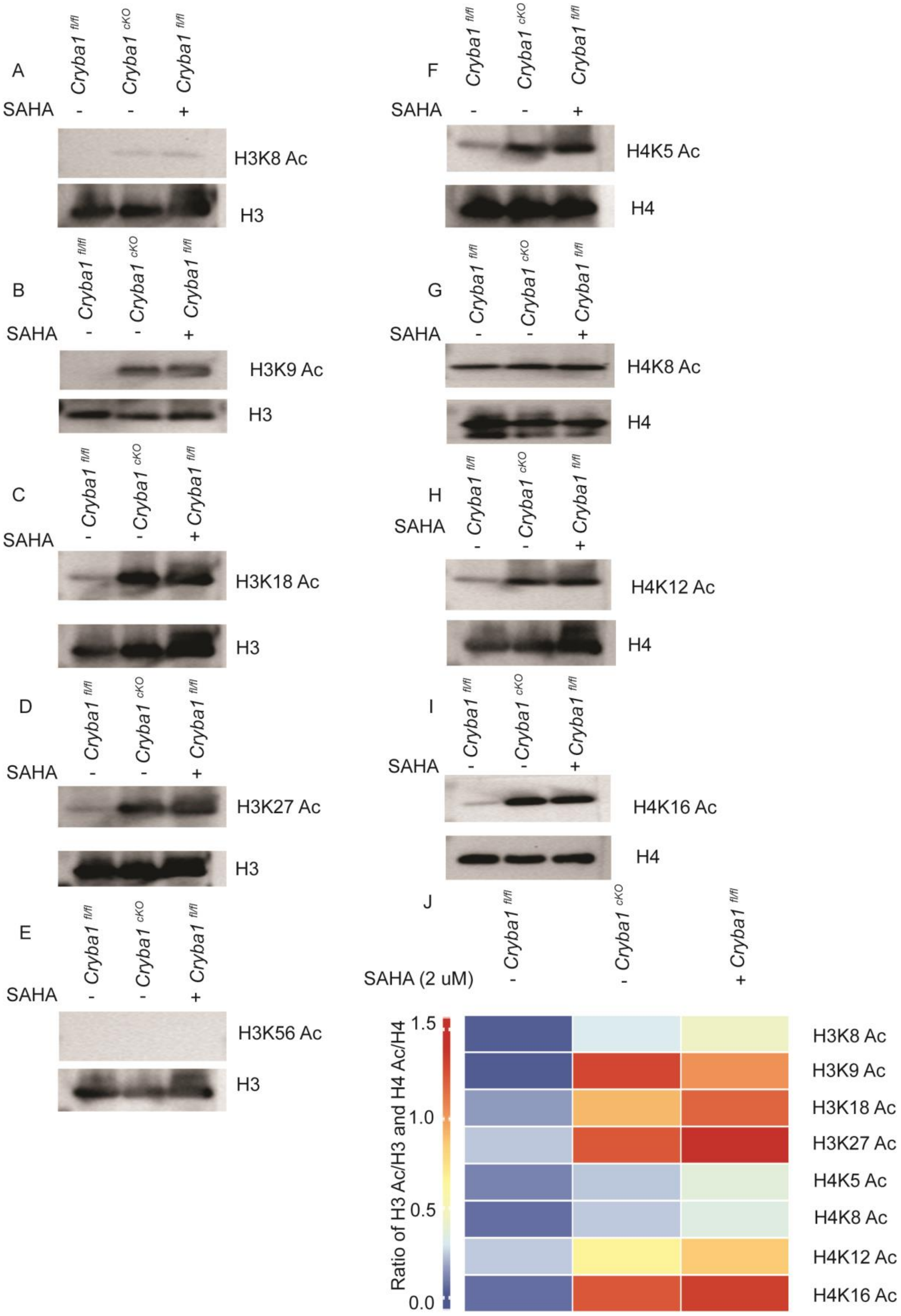


**<u>Extended Figure 2</u>: *Cryba1* deletion enhances histone acetylation in mouse RPE cells.** Western blot analysis demonstrated that histone acetylation increased in RPE cells from *Cryba1* cKO mice compared to *Cryba1*$^{fl/fl}$. *Cryba1*$^{fl/fl}$ RPE cells treated with 2 μM of the pan HDAC inhibitor SAHA was used as a positive control. Immunoblotting was performed against acetylated **(A)** anti-H3K8, **(B)** anti-H3K9, **(C)** anti-H3K18, **(D)** anti-H3K27, **(E)** anti-H3K56, **(F)** anti-H4K5 and **(G)** anti-H4K8 **(H)** anti-H4K12 and **(I)** anti-H4K16 followed by stripping and reprobing against anti-H3 or anti-H4 antibodies, respectively, followed by densitometric analysis **(J)**. Results are representative of three individual experiments. (n=3).

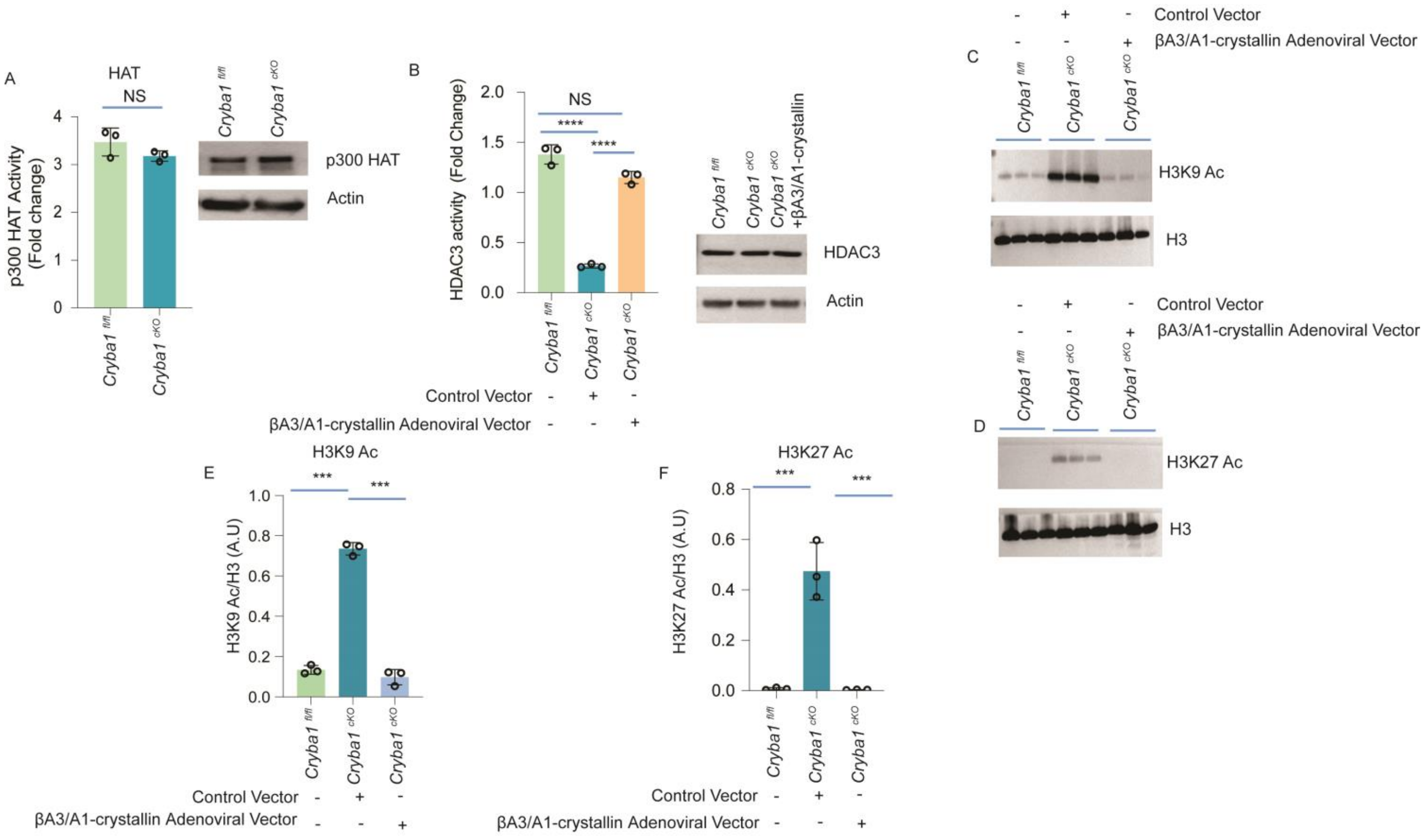


**<u>Extended Figure 3</u>: Overexpression of *Cryba1* can rescue HDAC3 activity and H3K9/27 acetylation in RPE cells. (A)** p300 activity in *Cryba1* cKO was comparable to *Cryba1*$^{fl/fl}$. p300 protein was immuno-precipitated from *Cryba1*$^{fl/fl}$ and *Cryba1* cKO RPE cells, followed by an *in vitro* HAT activity assay. IgG was used as a negative control. Data has been presented as fold change compared with blank value. Immunoblot analysis of p300 from total lysate isolated from *Cryba1*$^{fl/fl}$ and *Cryba1* cKO RPE cells was represented as input control, while actin was used as loading control. The result was representative of three individual experiments (n=3, NS=not significant) **(B)** *Cryba1* cKO RPE cells were stably over-expressed with *Cryba1* plasmid by using an adenoviral transfection procedure (*Cryba1* cKO+*Cryba1*). HDAC3 activity was significantly rescued in *Cryba1* cKO+Ad-*Cryba1* compared to *Cryba1*$^{fl/fl}$*.* Lysates isolated from *Cryba1*$^{fl/fl}$, *Cryba1* cKO*, and Cryba1* cKO+Ad-*Cryba1* were immuno-precipitated with anti-HDAC3 antibody followed by an analysis of HDAC3 activity. Immunoblot analysis against anti-HDAC3 antibody from the lysate isolated from *Cryba1*$^{fl/fl}$, *Cryba1* cKO*, and Cryba1* cKO+Ad-*Cryba1* RPE cells was used as control while actin was used as loading control. Results were representative of three independent experiments (n=3, NS=not significant, ****p<0.0001). **(C-D)** *Cryba1* cKO RPE cells were stably overexpressed with a wild type *Cryba1* adenoviral construct and vector control using an adenoviral transfection procedure. Immunoblot analysis indicated significant rescue in H3K9 and H3K27 acetylation in *Cryba1* cKO+Ad-*Cryba1.* Lysates isolated from *Cryba1*$^{fl/fl}$, *Cryba1* cKO*, and Cryba1* cKO+Ad-*Cryba1* RPE cells were immunoblotted with anti-H3K9-Ac **(C)** or anti-H3K27-Ac **(D)** antibodies. This was followed by stripping and reprobing against anti-H3 antibodies, which was used as a loading control. Results are representative of three individual experiments. (n=3). **(E and F)** represent densitometric analysis of H3k9/27 acetylation. n=3 (***p<0.001).

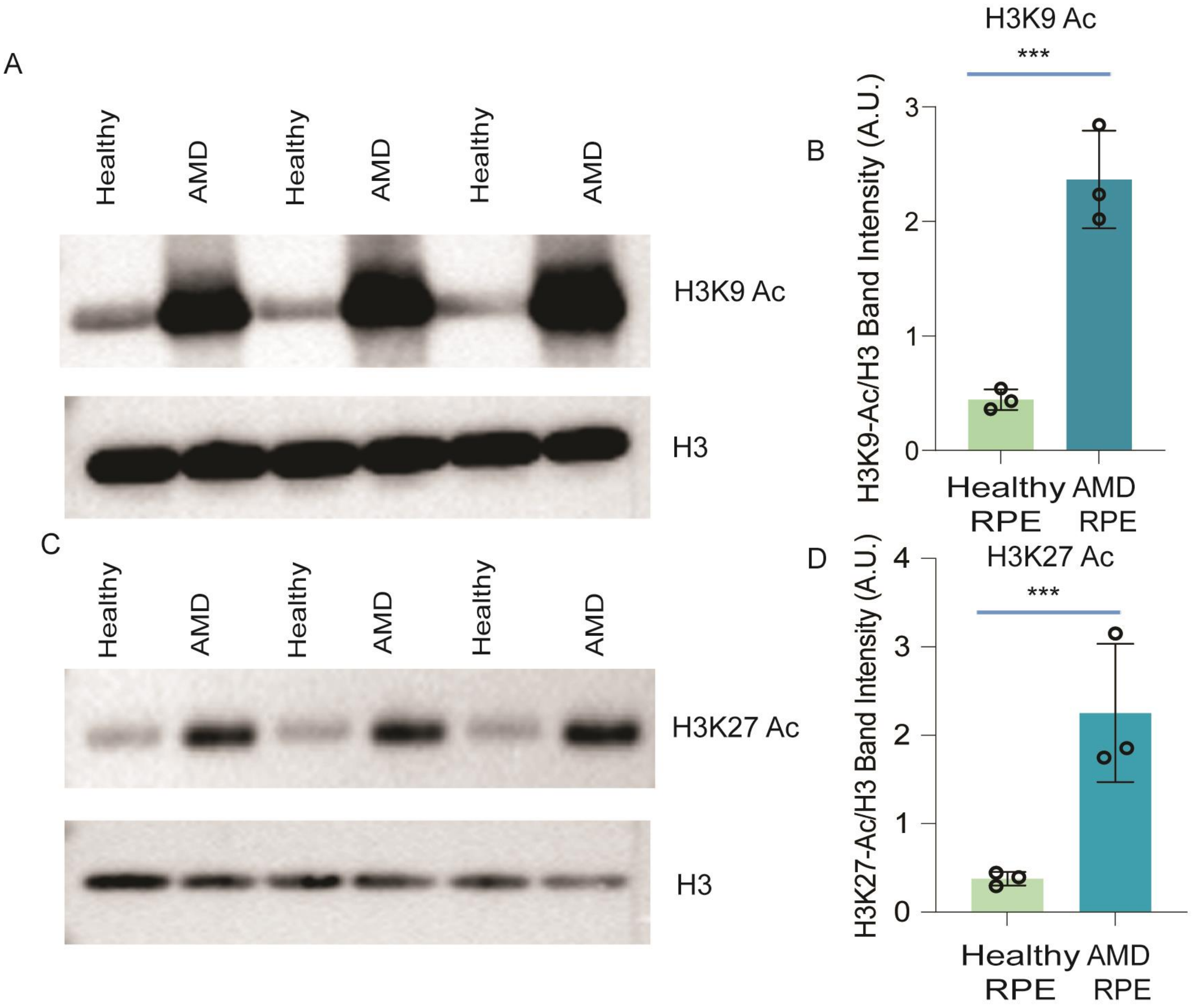


**<u>Extended Figure 4</u>: Increase in H3K9/H3K27 acetylation in human AMD. (A)** Immunoblot analysis indicated a significant increase in the level of H3K9 acetylation in AMD patients as compared to healthy human individuals*.* Lysates isolated from RPE cells of healthy and patients suffering from AMD were immunoblotted with anti-H3K9-Ac **(A)** or anti-H3K27-Ac **(C)** antibodies. This was followed by stripping and reprobing against anti-H3 antibody, which was used as a loading control (n=3). **B and D** represent densitometric analyses of H3k9/27 acetylation. Results are representative of three individual experiments. (n=3). n=3 (***p<0.001).

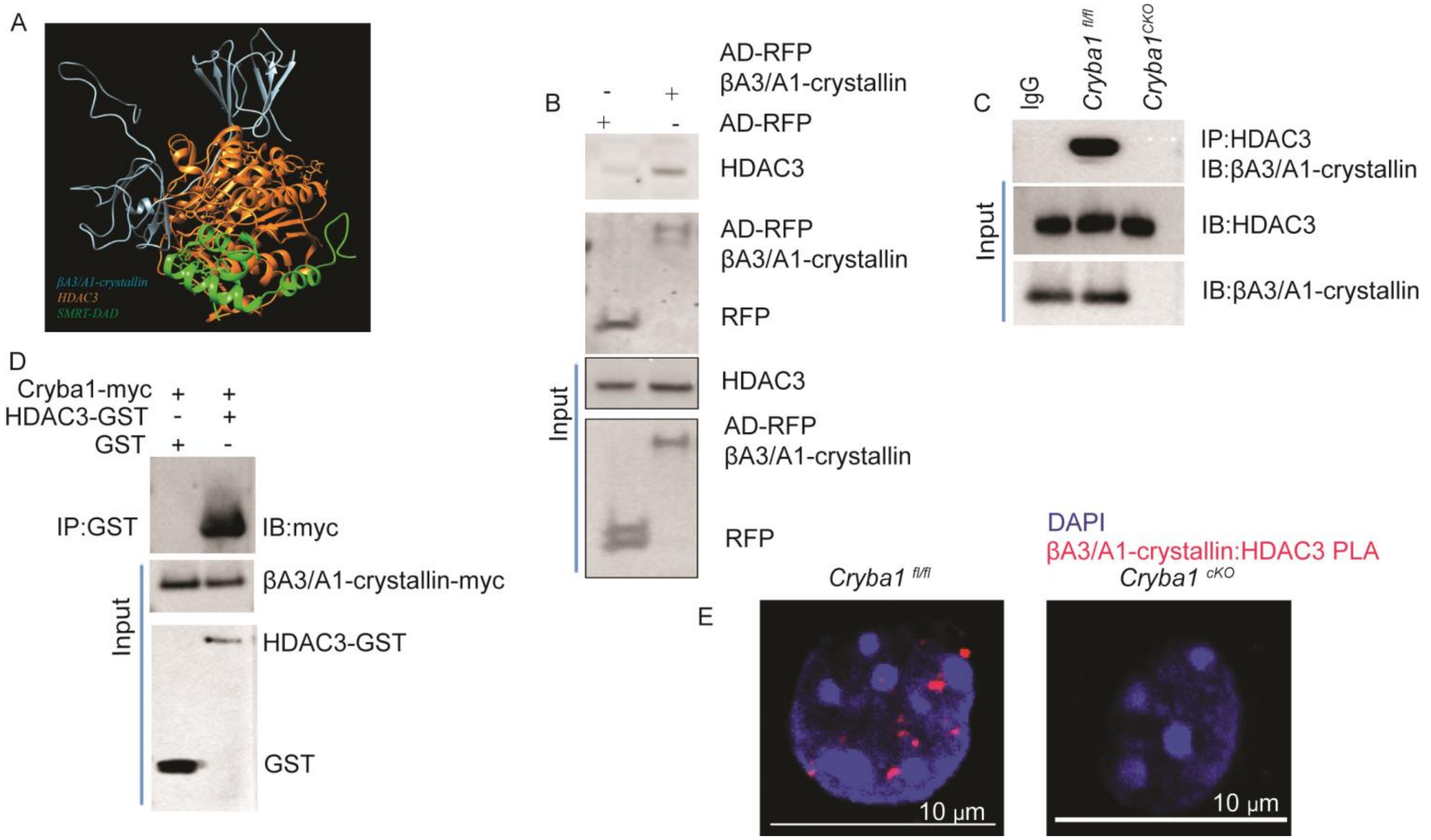


**<u>Extended Figure 5</u>: βA3/A1-crystallin physically interacts with HDAC3. (A)** *In silico* binding analysis predicted *Cryba1* binding with HDAC3 by using 3D computational modeling. βA3/A1-crystallin (blue), HDAC3 (orange), and *SMART-DAD* (green). **(B)** In the over expression system, βA3/A1-crystallin-RFP interacts with endogenous HDAC3. *Cryba1* cKO RPE cells were stably over-expressed with AD-RFP and βA3/A1-crystallin-RFP, respectively, followed by immunoprecipitation with anti-RFP antibody, then immunoblotted against anti-HDAC3 antibody. Whole cell lysates from the respective groups were then immunoblotted against anti-HDAC3 and anti-RFP antibodies, and were represented as input control (n=3). **(C)** Endogenous βA3/A1-crystallin binds to endogenous HDAC3. To analyze endogenous binding between HDAC3 and βA3/A1-crystallin, cell lysates isolated from *Cryba1*$^{fl/fl}$ and *Cryba1* cKO RPE were immuno-precipitated with anti-HDAC3 antibody followed by immunoblotting against βA3/A1-crystallin. IgG was used as a negative control. Immunoblotting against anti-HDAC3 and anti-*Cryba1* antibodies from the lysates isolated from *Cryba1*$^{fl/fl}$ and *Cryba1* cKO was used as input control (n=3). **(D)** Recombinant βA3/A1-crystallin-myc binds with recombinant HDAC3-GST in *in vitro* conditions. βA3/A1-crystallin-myc was incubated with either HDAC3-GST or

GST recombinant protein and then immunoprecipitated with anti-GST beads followed by immunoblotting against anti-Myc antibody. Whole cells lysates were immunoblotted against anti-myc, anti-HDAC3 and anti-GST antibodies (input control) (n=3). Results are representative of three individual experiments. (n=3). **(E)** The representative image illustrates the interaction between βA3/A1-crystallin and HDAC3 within the nucleus of the RPE in flat mounts. The PLA reveals prominent red puncta in the *Cryba1*$^{fl/fl}$ controls, indicating a robust interaction. Conversely, no puncta are observed in the nucleus of the *Cryba1* cKO RPE flat mounts, owing to the absence of crystallin. n=3, Scale bar=10μm.

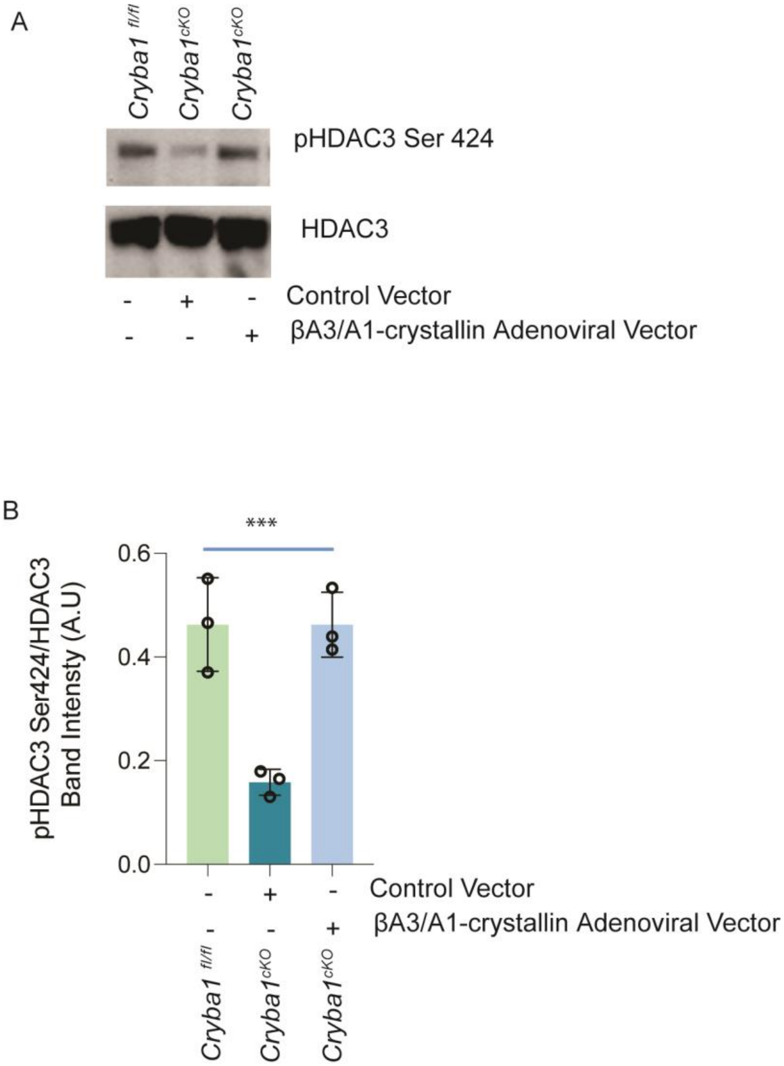


**<u>Extended Figure 6</u>: Overexpression of *Cryba1* can rescue HDAC3 phosphorylation at Ser424 in the *Cryba1* cKO RPE. (A)** Adenoviral-mediated overexpression of *Cryba1* significantly rescued HDAC3 phosphorylation at Serine 424. *Cryba1* cKO RPE cells were stably over-expressed with wild-type *Cryba1* construct or vector control using an adenoviral transfection procedure. Lysates isolated from *Cryba1*$^{fl/fl}$, *Cryba1* cKO+*vector control,* and *Cryba1* cKO+*Cryba1* were immunoblotted with anti-phospho HDAC3 Ser 424 antibody, followed by stripping and reprobing against an anti-HDAC3 antibody, which was used as a loading control. **(B)** Densitometric analysis of pHDAC3 Ser424/HDAC3 acetylation confirms the rescue of HDAC3 phosphorylation at Serine 424. Results are representative of three individual experiments. (n=3, ***p<0.001).

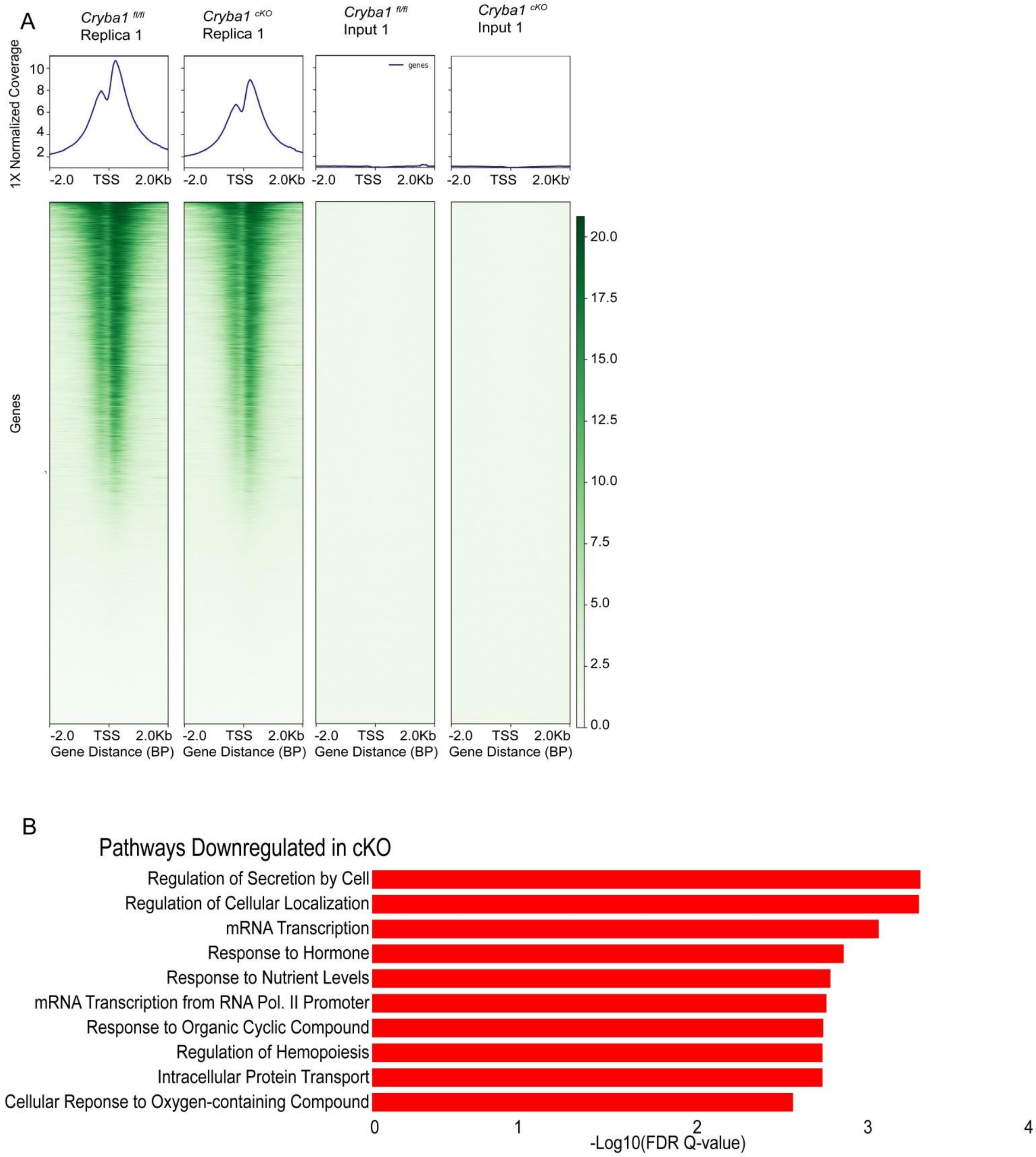


**<u>Extended Figure 7</u>: Deletion of *Cryba1* in RPE cells influences epigenetic changes. (A)** Heatmaps of chromatin occupancy of the histone marks H3K27Ac in mouse RPE cells. **(B)** Gene Ontology (GO) enrichment analysis depicting downregulated pathways in *Cryba1* cKO mouse RPE with reduced enrichment H3K27 acetylation peaks.
